# Aminoacyl tRNA Synthetases as Malarial Drug Targets: A Comparative Bioinformatics Study

**DOI:** 10.1101/440891

**Authors:** Dorothy Wavinya Nyamai, Özlem Tastan Bishop

## Abstract

Treatment of parasitic diseases has been challenging due to the development of drug resistance by parasites, and thus there is need to identify new class of drugs and drug targets. Protein translation is important for survival of plasmodium and the pathway is present in all the life cycle stages of the plasmodium parasite. Aminoacyl tRNA synthetases are primary enzymes in protein translation as they catalyse the first reaction where an amino acid is added to the cognate tRNA. Currently, there is limited research on comparative studies of aminoacyl tRNA synthetases as potential drug targets. The aim of this study is to understand differences between plasmodium and human aminoacyl tRNA synthetases through bioinformatics analysis. *Plasmodium falciparum*, *P. fragile*, *P. vivax*, *P. ovale*, *P. knowlesi*, *P. bergei*, *P. malariae* and human aminoacyl tRNA synthetase sequences were retrieved from UniProt database and grouped into 20 families based on amino acid specificity. Despite functional and structural conservation, multiple sequence analysis, motif discovery, pairwise sequence identity calculations and molecular phylogenetic analysis showed striking differences between parasite and human proteins. Prediction of alternate binding sites revealed potential druggable sites in PfArgRS, PfMetRS and PfProRS at regions that were weakly conserved when compared to the human homologues. These differences provide a basis for further exploration of plasmodium aminoacyl tRNA synthetases as potential drug targets.

## Introduction

Parasitic diseases like trypanosomiasis, malaria, leishmaniasis and filariasis affect millions of people in the world yearly [1–4]. These diseases cause a remarkable burden in economic development and health of affected countries and thus the need to come up with control and prevention strategies. Currently, the main mode of prevention and treatment of these parasitic diseases is by use of drugs as there are no approved vaccines in the market [5]. However, most parasites have developed resistance against conventional drugs leading to the drugs being ineffective [6,7]. Thus, there is need to develop new classes of drugs and to identify drug targets to solve the shortcoming of drug resistance. Targeting housekeeping pathways such as protein translation may help deal with drug resistance as they are important for the survival of most parasites [8–10].

Plasmodium parasite causes malaria disease, which is a major public concern due to its high mortality and morbidity rates [10,11]. There are five plasmodium species that cause malaria in human, and these are *Plasmodium falciparum* (*P. falciparum*), *Plasmodium vivax (P. vivax)*, *Plasmodium knowlesi* (*P. knowlesi)*, *Plasmodium malariae* (*P. malariae)* and *Plasmodium ovale* (*P. ovale*) [12]. Plasmodium has three genomes; cytoplasm, mitochondrial and apicoplast, and each of them needs a functional protein translation mechanism for growth and survival [10,13,14]. Plasmodium proteins involved in protein translation machinery are generally encoded by the nuclear genome and exported to target organelles to carry out various functions in protein synthesis [13,15–17].

Aminoacyl tRNA synthetases (aaRSs) are a group of key enzymes in protein translation pathway; they catalyze the first reaction, where an amino acid is added to the cognate tRNA molecule in the presence of ATP and magnesium (Mg^2+^) ions. This reaction takes place in two steps; first ATP activates the amino acid through formation of aminoacyl-adenylate intermediate, while the second step involves ligation of the adenylate intermediate to the cognate tRNA molecule through a covalent bond generating AMP [8,9,18]. Although the canonical function of these enzymes is to add amino acids to tRNA for translation and they are highly conserved in their catalytic domains, in general aaRSs show sequence, structural and functional diversity across organisms [19]. Furthermore, in some organisms, aaRSs have evolved to perform non-canonical functions such as angiogenesis, RNA splicing, signaling events, transcription regulation, apoptosis and immune responses [20–22]. *P. falciparum* tyrosyl-tRNA synthetases (PfTyrRS), for instance, have cytokine-like functions, while eukaryotic methionyl-tRNA synthetases (MetRS) have glutathione-S-transferase domains that play a key role in protein-protein interactions [23,24]. *P. falciparum* lysyl tRNA synthetase (PfLysRS) synthesizes diadenosine polyphosphate, a signaling molecule that plays a role in gene expression, DNA replication and regulation of ion channels of the parasite [25,26].

Of the five human malaria parasites, *P. falciparum*, known to be highly pathogenic, causes the most severe forms of malaria, and is responsible for most of the malaria mortality cases reported across the world [27]. *P. falciparum* has a total of 36 aaRSs that are asymmetrically distributed in either the cytoplasm, mitochondria or the apicoplast compartments. Of the 36 *P. falciparum* aaRSs, 15 reside in the apicoplast, 16 in the cytoplasm and four in mitochondria: AlaRS, GlyRS, ThrRS and CysRS are found both in the apicoplast and the cytoplasm and each of the four is encoded by a single gene and exported to the two compartments while only phenylalanine aminoacyl synthetase (PheRS) is encoded in the mitochondria [28–30]. *P. falciparum* protein translation in the mitochondria relies on enzymes imported from the cytoplasm including aaRSs [28]. The apicoplast encodes AspRS, PheRS, ValRS, LysRS, HisRS, AsnRS, ProRS, SerRS, TrpRS, ArgRS, IleRS, GluRS, LeuRS, TyrRS and MetRS while AlaRS, CysRS ThrRS and GlyRS are reported to have a single gene encoding both the cytoplasm and apicoplast enzyme [15,17,30,31]. A single transcript for each gene is spliced alternatively to generate the two isoforms for each protein which are then targeted to either the cytosol or the apicoplast [29,30]. Each of these genes encodes a protein with a N-terminal extension that corresponds to a signal and transit peptide and is conserved in the apicomplexa phylum [29]. *P. falciparum* cytoplasm has genes that encode ProRS, AspRS, IleRS, LysRS, HisRS, PheRS, AsnRS, ArgRS, GlnRS, SerRS, TrpRS, ValRS, MetRS, LeuRS, GluRS and TyrRS [31].

In human, aaRSs carry out aminoacylation reactions in the cytoplasm, nucleus and the mitochondria. After tRNA is encoded in the nucleus, it is transported to the cytoplasm where protein translation takes place [8]. The human mitochondria acquires nuclear-encoded aaRSs with the aid of translation signals within the aaRSs proteins to carry out protein synthesis [32]. The cytoplasm is the only compartment where both aminoacylation and protein synthesis exclusively takes place in humans. Human aaRSs are, thus, classified as mitochondrial or cytoplasmic based on the compartment where they are localized [32]. In human, a total of 36 aaRSs have been reported with 17 of them in the mitochondrion and 16 aaRSs exclusively functioning in the cytoplasm while the other three catalyze aminoacylation reactions in both organelles [8,32]. The three bifunctional aaRSs in human are GlnRS, GlyRS and LysRS. In the cytoplasm, aminoacylation of proline and glutamate is catalyzed by a single bifunctional enzyme (Glu/ProRS). Thus, both compartments have enzymes for charging all the 20 amino acids [32,33].

Generally, aaRSs proteins are classified into two distinct classes based on key features of the catalytic site architecture and the manner of charging tRNA [18,20]. Class I aaRSs include IleRS, LeuRS, MetRS, CysRS, GlnRS, GluRS, TrpRS, ValRS, ArgRS and TyrRS. Proteins in this class have a catalytic domain (Figure 1A) characterized by a Rossmann fold (RF) located near the N-terminal [34]. The catalytic domain of this class comprises five parallel β-sheet strands flanked by α-helices. The RF possesses highly conserved HIGH and KMSKS motifs separated by a loop [35,36] as shown in Figure 1A. The HIGH motif is located in a region formed by a loop linking the first β-sheet strand and the adjacent α-helix while the KMSKS motif occurs after the fifth β-sheet strand [8]. The RF domain has an insert known as the connective peptide I (CPI) in all enzymes in this class whose structure is characteristic of mixed α and β folds. Proteins in this class have common domains that include an alpha-helical anticodon binding domain (ABD), connective peptide (CPI) and the tRNA stem contact fold [37]. The CPI insert is found towards the end of the first half of the fourth β-strand of the RF joining the N-terminal and C-terminal sections of the catalytic domain [8]. With the exception of TyrRS, MetRS and TrpRS all Class I enzymes are monomeric [8]. In monomeric enzymes, the CPI binds tRNA at the 3’-single stranded end while in TrpRS and TyrRS it forms the dimer interface of these dimeric enzymes [34,38]. In ValRS, IleRS and LeuRS, the CPI insert is enlarged (250-275 amino acid residues as compared to CysRS and MetRS where it is 50 and 100 residues respectively) to include an editing domain for editing misacylated tRNA through hydrolysis [39]. The editing domain proofreads the aminoacylation process through pre-transfer or post-transfer editing [8]. Post-transfer editing involves hydrolyzing of misacylated tRNA to amino acid and tRNA while pre-transfer modification hydrolyzes the mis-activated aminoacyl adenylate to AMP and amino acid [8]. The ABD of proteins in Class I occurs at the C-terminal which binds the anticodons in the cognate tRNA [40].

**Figure 1.**
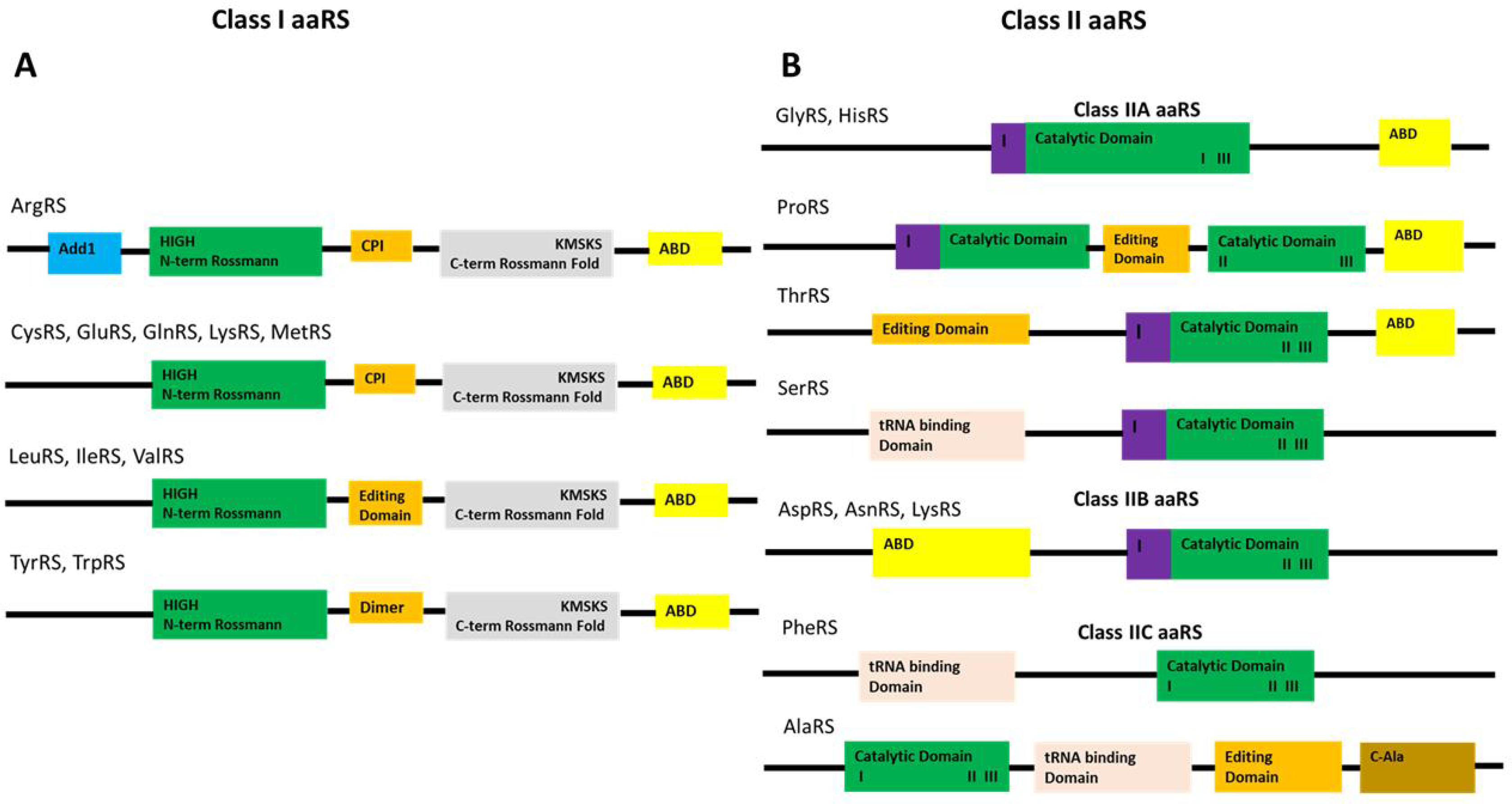
Key domains of aminoacyl tRNA synthetases. **A)** Class I aaRS showing the Catalytic Domain (CD) and the anticodon binding domain (ABD). The CD has a CPI insert in all the proteins in this Class. The CPI insert (orange) in IleRS, LeuRS and ValRS is enlarged to form an editing domain while in TyrRS and TrpRS it functions in the formation of dimers. ArgRS has an Add1 domain (cyan) at the N-terminus which is involved in tRNA recognition. **B)** Class II aaRS showing the catalytic domain with the three conserved motifs (I, II and III). In GlyRS, HisRS and ProRS the anticodon binding domain is at the C-terminal. In AspRS, AsnRS and LysRS it is at the N-terminal, while in AspRS, LysRS and AsnRS proteins the ABD occurs at the C-terminal. Dimer interfaces are shown by a magenta color and are characterized by motif I. ProRS has an editing domain that occurs between motif I and II at the catalytic site while in ThrRS the editing domain is located at the N-terminal. AlaRS has a C-Ala domain (gold) at the C-terminal that functions in dimer formation.

Class I enzymes binds to the tRNA acceptor end through the minor groove and these enzymes aminoacylate the 2’-OH group of adenosine nucleotide [8,40]. Proteins in this class can further be classified into five subclasses based on sequence similarity and physicochemical properties of their substrates [41,42]. Subclass Ia members charge hydrophobic amino acids that have aliphatic side chains and include ValRS, MetRS, IleRS and LeuRS. Subclass Ib proteins have charged amino acids as their substrates and include GlnRS, CysRS and GluRS. Members of subclass IIb bind to the cognate tRNA before carrying out the aminoacylation process [8,43]. TrpRS and TyrRS belong to subclass Ic and their substrates are aromatic amino acids. ArgRS is the only member of subclass Id and it possesses an Add1 domain at the N-terminal whose function is to recognize the D-loop in the tRNA core (Figure 1A) [8,40]. Class I LysRS found in some bacteria and archaea shares structural similarity with subclass Ib but it has a unique alpha helix cage and is thus grouped in subclass Ie [44].

Class II aaRSs include HisRS, ProRS, LysRS, SerRS, AspRS, ThrRS, AlaRS, GlyRS, PheRS and AsnRS. Proteins in this class are further grouped in three subclasses whose members are more closely related than other subclasses [45,46]. Class IIa proteins exist as dimers and includes ProRS, SerRS, GlyRS, ThrRS, HisRS and all have the aminoacylation domain at the N-terminal [8]. Members of this subclass have an ABD at the C-terminal (figure 1B). The anticodon binding domain is absent in SerRS as this protein does not require an anticodon to discriminate its cognate tRNA [47,48]. ProRS has editing domains located between motifs I and II at the catalytic domain while in ThrRS the editing domain is at the N-terminus (Figure 1B) [40]. Members of Class IIb are dimers and have a C-terminal catalytic domain that is structurally similar and include AspRS, LysRS and AsnRS. The ABD in this subclass is located at the N-terminal (Figure 1B). Class IIc includes PheRS, AlaRS and GlyRS and all exist in tetrameric conformation [8,46]. AlaRS possesses a C-Ala domain at the C-terminal which is absent in other members of Class IIc. The editing domain in AlaRS occurs between the tRNA binding domain and the C-Ala domain (Figure 1B) [40].

Class II enzymes possess a catalytic site domain characterized by seven β-sheet strands connected by α-helices [49]. This domain, just like the Class I catalytic domain couples amino acid, ATP and tRNA 3’-terminus during catalytic reactions [40,50]. Class II catalytic domain has three weakly conserved motifs (Figure 1B, Figure 2B); Motif I found at the N-terminal of the catalytic region is characterized by a long α-helix linked to a short β-strand with a proline residue at the end which is highly conserved and is involved in homo dimerization [51]. Motif II juxtaposes amino acid, ATP and tRNA and comprise β-sheet strands. Motif III is located at the C-terminal of the catalytic domain and binds ATP and comprise alternate β-strands and α-helices [36]. LysRS can be classified in both classes based on the structure and mode of charging tRNA, with Class I LysRS occurring in some bacteria and most archaea [52] while Class II LysRS occurs in most bacteria and all eukaryotes [8].

**Figure 2.**
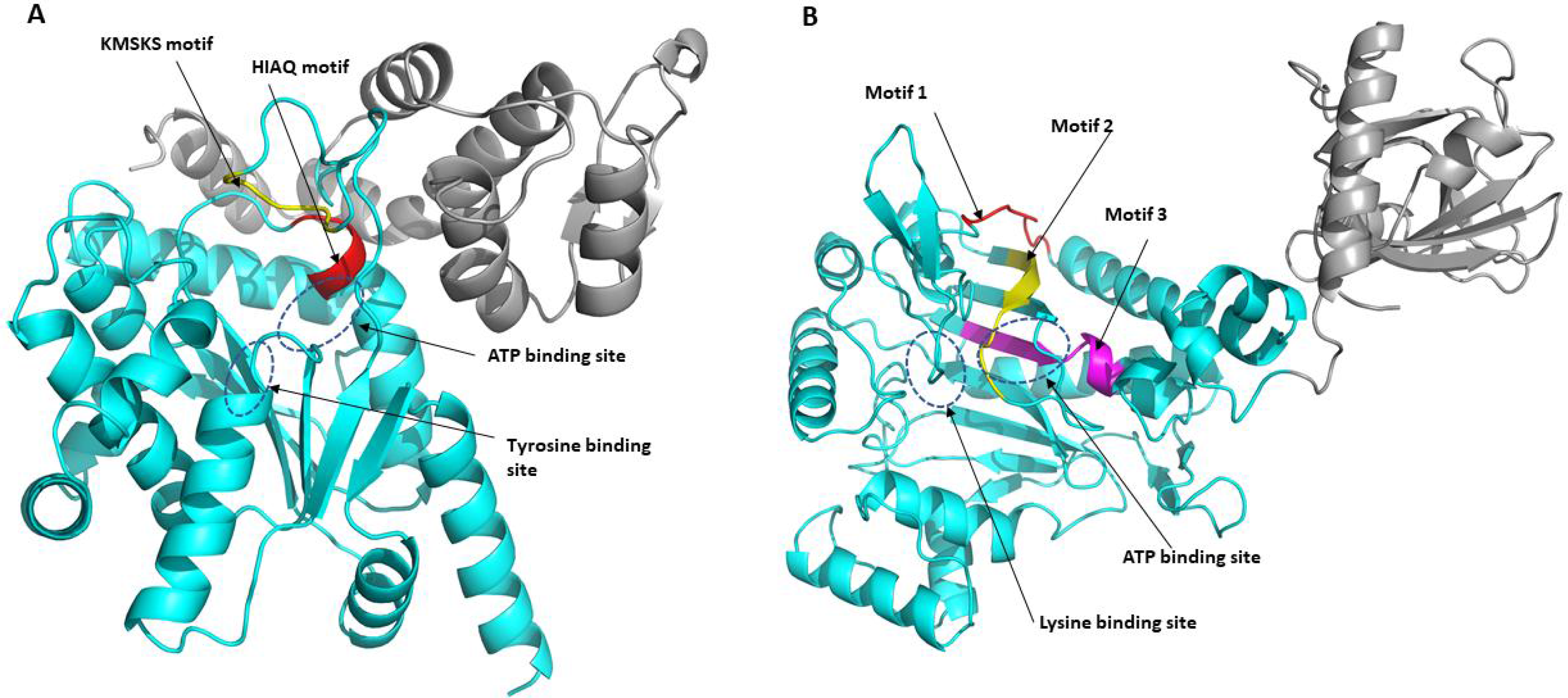
A) The apo structure of PfTyrRS (Class I) The catalytic domain (residues 22-260) is shown as cyan (cartoon) while the anticodon binding domain (residues 261-370) is shown in grey. The highly conserved KMSKS motif (red) and the HIGH motif (yellow) are shown in the structure. ATP and tyrosine binding sites are shown as blue dotted ellipses. Asp61, His70, Ala72, Gln73, Gln210, His235, Met237, Leu238, Met248, Lys250 are involved in ATP binding while residues Tyr60, Glu64, Ala96, Phe99, Ile172, Tyr188, Gln192 and Asp195 are involved in tyrosine binding [53]. B) **Apo structure of PfLysRS (Class II)**. The anticodon binding domain (residues 77-226) is shown in grey cartoon while the catalytic domain (residues 227-583) is shown in cyan. Motif I (red), motif II (yellow) and motif III (magenta) are shown by arrows. The ATP binding site and lysine binding site are shown by the blue dotted ellipses. Residues Arg330, His338, Asn339, Phe342, Glu500, Asn503, Gly556 and Arg559 are implicated in binding of ATP while residues Glu308, Asn330, Glu346, Tyr348, Asn503, Tyr505, and Glu507 are involved in binding of lysine [54].

Protein translation has been explored as a target in the development of antimalarial drugs with most drugs interfering with the ribosome [55]. Recently, there has been increased interest in exploring *P. falciparum* aaRSs as potential drug targets [15,25,31,55–57]. Plasmodium aaRSs inhibitors have been identified that target either the ATP pocket, the amino acid or tRNA binding site or the editing domains of some of these enzymes. Some of the compounds reported to target *P. falciparum* aaRSs are halofuginone, cladosporin, 3-aminomethyl benzoxaborole AN6426, glyburide and TCMDC-124506 [58–61]. Halofuginone, a derivative of febrifugine, targets ProRS tRNA and proline binding site mimicking tRNA 3’-Adenine 76 and L-pro in an ATP dependent manner [57,62,63]. Halofuginone binding to human and plasmodium ProRS involves identical residues and in both the compound mimics proline and adenine substrates binding pose thus leading to toxicity in human cells [58,64,65]. Cladosporin, a secondary metabolite from fungi, is reported to have activity against blood and liver stage *P. falciparum* and its activity is selective to only the parasite LysRS protein [25,59]. Cladosporin, an adenosine analogue binds at the ATP binding site of PfLysRS [25,59]. Cladosporin can, thus, be used as a basis for development of other scaffolds with improved drug-like properties. The compound 3-aminomethyl benzoxaborole AN6426 was reported to be active against LeuRS in drug resistant *P. falciparum* but did not impair growth of the wild type [61]. This compound binds to the editing domain of PfLeuRS and inhibits it inactivating the 3’Adenine 76 nucleotide of the cognate tRNA covalently and the catalytic turnover of *P. falciparum* resistant strains [56]. Glyburide and TCMDC-124506 are reported to bind to a site adjacent to the ATP binding site of PfProRS and displace key residues involved in ATP binding thus inhibiting the enzyme activity [60]. Glyburide and TCMDC are selective to PfProRS and do not cause toxicity to human cells and thus can be used as a basis for development of drugs targeting PfProRS.

Due to these shortcomings of the current compounds that target aaRSs and the ever-increasing antimalarial drug resistance [6,15,16,19,27,66], there is need to develop novel drugs and identify more targets to counter this resistance. In addition, the development of drugs that are active against the liver, blood stage parasites [66] and the sexual stages of the parasites thus terminating the infection cycle would help in malaria eradication [67]. With aaRSs proteins being present in all stages of the parasite life cycle, identification of subtle differences between the plasmodium and human proteins would help in achieving this goal.

Although aaRS are desirable drug targets, selectivity of drugs to only parasitic aaRS and not human proteins is a challenge as human aaRS have bacterial and eukaryotic origin [68–70]. High conservation of aaRS across plasmodium and the human host may hinder development of parasite specific inhibitors [10,71,72]. Comparative studies between host and parasite sequences and structures are important in identifying differences that can be exploited for drug development [71,73–75]. The aim of this study was to discern sequence and structural differences of aaRS between human and plasmodium proteins despite the functional conservation of these proteins. The differences that occur at the active pockets and the predicted druggable sites can thus be exploited for development of drugs with good selectivity [76]. This study included *P. falciparum, P. bergei*, *P. malariae*, *P. ovale*, *P. yoelii*, *P. vivax*, *P. knowlesi*, *P. fragile*, human and other mammalian sequences. Sequences from toxoplasma and cryptosporidium species which belong to the apicomplexa family and other prokaryotes were also included for molecular phylogenetic tree calculations (Additional file 1). Targeting of cytosolic protein machinery in plasmodium shows immediate death while inhibition of apicoplast protein translation machinery is reported to show delayed death where parasites die only during the next replication process after treatment [77]; thus, in this study we focus mainly on the cytosolic aaRSs. The sequences were classified into two groups based on differences in structure of their catalytic domain and further into the different aaRS families based on their amino acid substrates [18,20,78]. The study was divided into two parts. First, sequence-based analysis which involved motif search, multiple sequence alignment and phylogenetic tree calculations was carried out. Secondly, structure-based analysis was carried out which involved modeling of 3D structures of proteins, mapping of identified motifs to these structures and identification of probable allosteric drug targeting sites on the 3D models. The results showed striking differences in motifs and at residue level between parasite and human proteins. The results from this study thus form a basis for further research on aaRS as potential antimalarial drug targets and other parasitic diseases.

## Methods

### Sequence retrieval

*Plasmodium falciparum* aminoacyl tRNA synthetases (PfaaRS) were retrieved from NCBI-Protein database [79]. Protein sequences of other plasmodium species and human ones were searched by BLAST in UniProt using each PfaaRS as the query sequence for the specific family [80]. The BLASTp algorithm with the default BLOSUM62 matrix was used for the search of homologous sequences. (Additional file 1). The data set consisted of the five plasmodium species that infect human, *P. bergei*, *P. yoelii*, *P. fragile* and human homologues. For phylogenetic tree calculations, other apicomplexan (cryptosporidium and toxoplasma) sequences and prokaryote sequences were also retrieved (Additional file 1). The sequences were then grouped into 20 groups based on the different aaRS families. Retrieved sequences were also grouped into two classes (Class I and Class II), each consisting of ten protein families [81,82]. Crystal structures for human and *P. falciparum* ArgRS, TrpRS, MetRS, TyrRS, LysRS and ProRS proteins were retrieved from Protein Data Bank (PDB) [83].

### Motif discovery

Motif discovery was done using Multiple Expectation Maximisation for Motif Elicitation (MEME) vs 4.11 to identify highly conserved motifs in each aaRS class [84]. A total of 90 motifs with a motif width of 6-50 residues were run for each of the non-homologous classes. The MAST tool was used to identify overlapping motifs [85]. A Python script was used to analyse MAST files and MEME log files. Motif conservation was represented as a number of sites per a total number of class sequences, and the results were displayed as heatmaps. Further, motif discovery was performed for each aaRS family and the results also displayed as heatmaps. For each aaRS family, the default parameters were used with motif width of 6-50 residues and the number of motifs run for each family varied (Additional file 3).

### Homology modelling and model quality assessment

3D structures of *P. falciparum*, *H. sapiens, P. bergei*, *P. malariae*, *P. knowlesi*, *P. fragile*, *P. yoelii*, *P. ovale* and *P. vivax* proteins were built by homology modelling using MODELLER v9.15 [86]. Templates were identified using HHpred and PRotein Interactive MOdeling (PRIMO) webservers [87,88] for the six ArgRS, TyrRS, TrpRS, MetRS, LysRS and ProRS families (Additional file 2). The other families had no good quality templates hence models were not built. For ArgRS - 5JLD [89]; for MetRS - 4DLP [90]; for TrpRS - 4J75 [91]; for TyrRS - 5USF [92]; for ProRS - 4NCX [93] and for LysRS - 4DPG [94] was used. For each protein, 100 models were calculated and the top three models with the lowest z-DOPE (Discrete Optimized Protein Energy) score were selected for validation. Structure quality assessment was done using Protein Structure Analysis (PROSA) webserver [95], Verify3D [96] and Qualitative Model Energy Analysis (QMEAN) [97] and the model with the best scores was selected for allosteric site prediction and motif mapping.

### Sequence alignment

For each family of sequences, multiple sequence alignment was carried out using Profile Multiple Alignment with Local Structures and 3D constraints (PROMALS3D) and Tree-based Consistency Objective Function Evaluation (TCOFFEE) alignment tools [98,99]. Visualization and editing of the alignments were done using the Jalview vs. 2.10 software [100]. The alignment results from the two alignment tools were compared, and, in both, it was observed that the sequences were aligned identically except for the less conserved C-terminal and N-terminal regions. TCOFFEE sequence alignments were used for the phylogenetic tree calculations as well as for all versus all pairwise sequence identity calculations via a Python script. The sequence identity results were translated into heatmaps using a Matlab script.

### Molecular phylogenetic analysis

Phylogenetic tree calculations were carried out for each family of aaRSs to study evolutionary relationships within the protein families using Molecular Evolutionary Genetic Analysis (MEGA) vs7.0 tool [101]. For sequence alignment of each family, three gap deletion options - 90%, 95% and 100% - were used to calculate the models, and the best three models for each deletion option were selected based on the lowest Bayesian information criterion (BIC) scores. Maximum Likelihood (ML) statistical method was used to infer evolutionary relationship while calculating trees for the top three models for each gap deletion option for each protein families [102]. Total of 180 (3×3×20) trees were calculated. Nearest-Neighbor-Interchange search was performed for all the constructed trees. BioNJ and Neighbor Join algorithms were used for a matrix of pairwise distances calculated using JTT model to obtain the initial trees for the heuristic search and the topology with the highest log likelihood selected [103]. A strong branch swap filter and 1000 bootstrap replicates were used for each tree calculation. The trees were then compared to the bootstrap consensus trees to ensure that branching patterns were accurate and the best model and gap deletion for each case was, then, chosen.

### Prediction of alternate druggable sites

Structure-based drug design and development requires understanding of the structure and function of the binding sites of the target protein. Identification of new drug targeting sites different from the validated active sites is key in development of new classes of drugs. In this study, probable druggable sites of our protein models were determined using FTMap webserver [104] and SiteMap [105,106]. Homology models were used as input for the prediction of probable druggable sites. The FTMap webserver identifies probable binding sites by screening of small compounds that vary in shape, polarity and size using an empirical energy function and the CHARMM force field [104]. The webserver docks isopropanol, acetaldehyde, phenol, benzaldehyde, urea, dimethyl ether, acetonitrile, ethane, acetamide, benzene, methylamine, cyclohexane, ethanol, N,N-dimethylformamide, isobutanol and acetone at the surface of the protein [107]. Clusters of low energy conformations are calculated and ranking of the probes is done based on the average energy [104]. The site that binds most of the compounds is considered the active binding site while other regions that bind several compounds are the predicted binding sites.

SiteMap, a tool in Schrödinger suites assigns site points in cavities that are likely to contribute to protein-protein or protein-ligand interactions based on energetic and geometric properties [105,106]. The tool uses an algorithm that depends on how well sheltered the sites from solvents are and how close they are from the protein surface to determine the likeness of a site point. The sites are classified based on different properties which include; how enclosed is the site by the protein, the size of the site as measured by the number of points, the degree by which a ligand can accept or donate hydrogen bonds, how tight the site interacts with the protein, how exposed the site is to the solvent and the hydrophilic and hydrophobic nature of the site [105]. The predicted binding sites are then ranked based on a SiteScore calculated using a linear combination of these factors [105].

## Results and Discussion

In this study, 92 Class I and 89 Class II proteins were analysed for the eight plasmodium species and their human homologues. More mammalian sequences were included for MSA and motif search within each aaRS family to avoid bias. A protein from each aaRS family was represented for each organism except for PbAspRS which was reported as a putative protein thus we did not include it in the study. Overall, the study is divided into two parts. In the first part, sequence related analyses such as MSA, phylogenetic tree calculations and motif identification were performed with the aim of understanding the general differences between plasmodial and human proteins. The second part included homology modelling, mapping of motif information into 3D structures and identification of alternative drug targeting sites, as the active site within a family of proteins is generally highly conserved, hence identification of plasmodial protein specific inhibitors might be challenging.

### PART 1 - SEQUENCE BASED ANALYSES

#### Discovery of motifs that are conserved in each AARS class

Motif analysis was done for each aaRS class (Figure 3 and 4) and for each family (see Additional file 3 & 4). The results were displayed as heatmaps using a Python script and mapped to multiple sequence alignment results and available structures. Motifs discovered for each family varied as shown in Additional file 3 and 4. Motif numbering used in this section is based on the MEME results.

**Figure 3.**
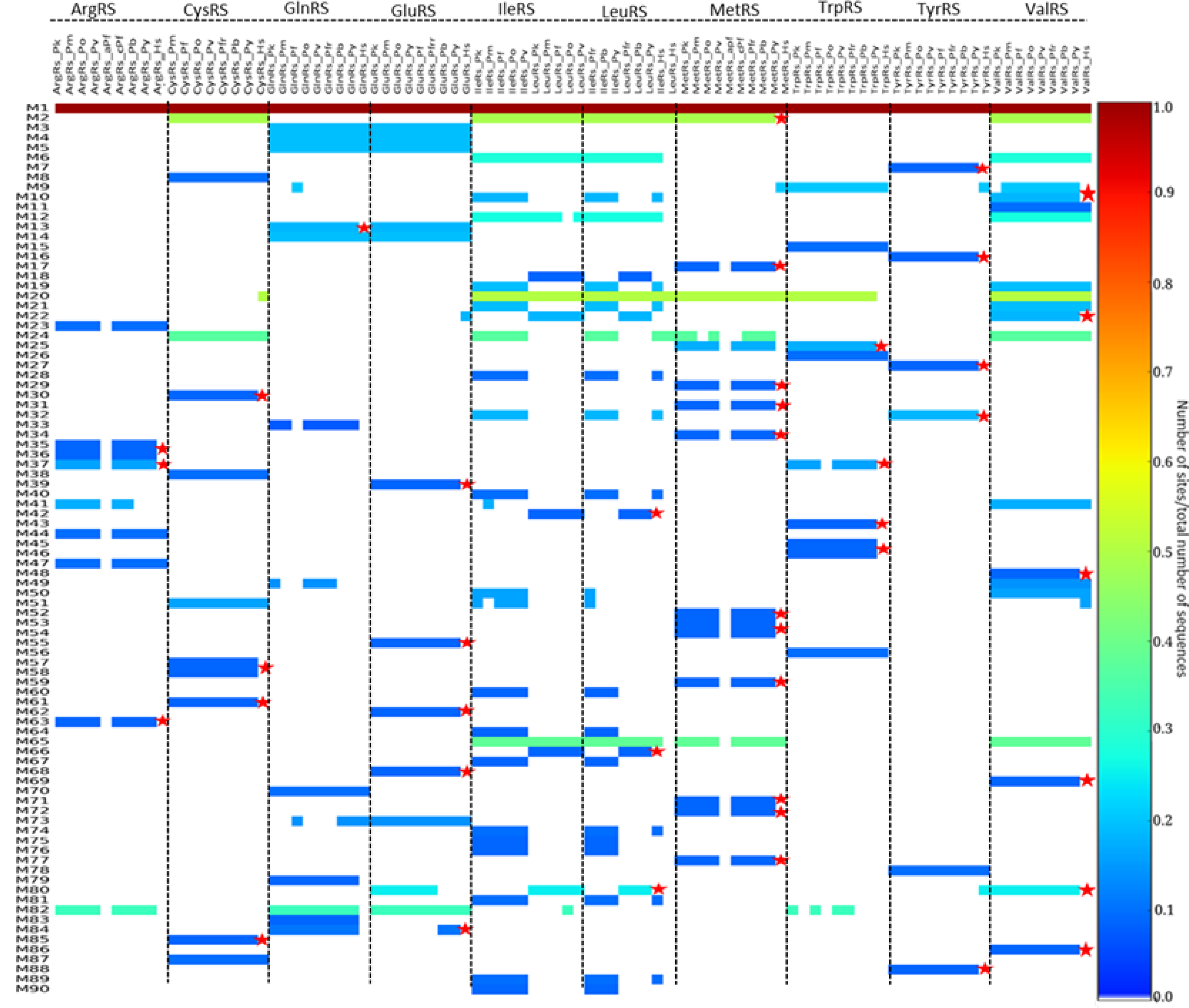
Motifs identified in Class I aaRS presented as a heat map. The colours represent conservation of motifs of the identified 90 motifs in this class. Conservation increases from blue to red while the absence of motifs is shown by a white colour. Motifs not present in human aaRS are shown in a red asterisk.

**Figure 4.**
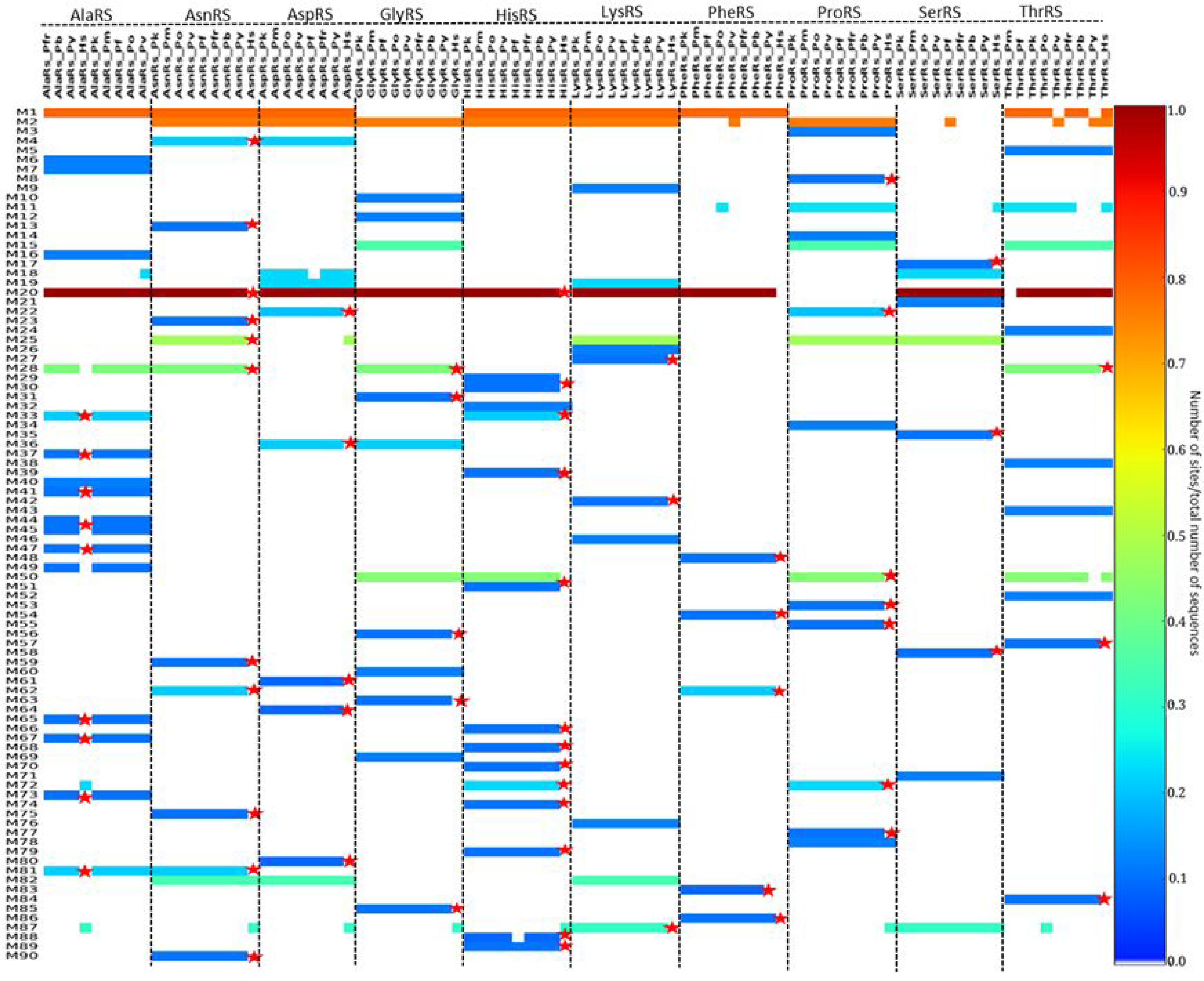
Motifs identified in Class II aaRS presented as a heat map. Motifs not present in human aaRS are shown in a red asterisk. The colours represent conservation of the identified 90 motifs in this class. Conservation increases from blue to red while the absence of motifs is shown by a white colour.

In Class I, 90 motifs were identified as shown in Figure 3. The start and end positions of highly conserved motifs in this class is shown in Table 1. Motif 1 was conserved in all 92 sequences in this class (Figure 3). This motif contains conserved residues involved in ATP binding. Motif 2 was present in 45 out of 92 sequences and this motif has also been reported to be important in ATP binding [40]. Class I aaRS enzymes are known to have a Rossmann fold catalytic domain which is characteristic of the highly conserved Motif 1 and 2 [108,109]. Motif 12, 20 and 65 were also highly conserved among sequences in this class. The other motifs clustered based on the enzyme family but some were conserved across different enzymes within the same class. Motif 3, 4, 5, 13 and 14, for example, was conserved in all GluRS and GlnRS sequences (Figure 3). These shared motifs show that these two proteins have a high sequence identity and may explain why plasmodium apicoplast GluRS mischarges glutamine specific tRNA with glutamate. In this case, glutamate is then changed to glutamine a reaction catalysed by glutamyl-tRNA amidotransferase enzyme [110,111].

**Table 1.** Highly conserved motifs in Class I aaRS. Motif 1, 2, 6, 12, 20 and 65 in Class I *P. falciparum* aaRS and the human homologues. Motif positions in the sequences are indicated and dashes are used where the motif is not present.

|  | Motif 1 | Motif 2 | Motif 6 | Motif 12 | Motif 20 | Motif 65 |
| --- | --- | --- | --- | --- | --- | --- |
| PfArgRS | 134-152 | ---- | ---- | ---- | ---- | ---- |
| HsArgRS | 197-215 | ---- | ---- | ---- | ---- | ---- |
| PfCysRS | 131-149 | 382-419 | ---- | ---- | ---- | ---- |
| HsCysRS | 53-71 | 405-442 | ---- | ---- | 700-714 | ---- |
| PfGlnRS | 284-302 | ---- | ---- | ---- | ---- | ---- |
| HsGlnRS | 266-284 | ---- | ---- | ---- | ---- | ---- |
| PfGluRS | 313-331 | ---- | ---- | ---- | ---- | ---- |
| HsGluRS | 200-218 | ---- | ---- | ---- | ---- | ---- |
| PfIleRS | 127-145 | 782-819 | 623-651 | 222-271 | 167-181 | 91-111 |
| HsIleRS | 44-62 | 599-636 | 444-472 | 139-188 | 94-108 | 63-83 |
| PfLeuRS | 141-159 | 1119-1156 | 688-716 | 219-268 | 181-195 | 160-180 |
| HsLeuRS | 49-67 | 715-752 | ---- | ---- | 89-103 | ---- |
| PfMetRS | 228-246 | 514-551 | ---- | ---- | 268-282 | 247-267 |
| HsMetRS | 269-287 | ---- | ---- | ---- | 310-324 | 289-309 |
| PfTrpRS | 306-324 | ---- | ---- | ---- | 534-548 | ---- |
| HsTrpRS | 159-177 | ---- | ---- | ---- | ---- | ---- |
| PfTyrRS | 61-79 | ---- | ---- | ---- | ---- | ---- |
| HsTyrRS | 53-71 | ---- | ---- | ---- | ---- | ---- |
| PfValRS | 86-104 | 628-665 | 454-482 | 181-230 | 126-140 | 105-125 |
| HsValRS | 340-358 | 861-898 | 708-736 | 435-484 | 380-394 | 359-379 |

Motif 1 consisting of the HIGH signature which is characteristic of the Rossman fold was conserved in all Class I aaRS (Figure 3) [81]. This class also showed high conservation of a Motif 2 containing the KMSKS conserved signature which has also been reported to be part of the RF in this class (Figure 6 & Additional file 3 & 4). The HIGH motif is present in the first half of the RF while the KMSKS motif is present in the second half of the RF domain (Figure 5 & Additional file 4). Motif conservation of the Rossman fold reflects the functional importance of this region. This fold is involved in ATP binding and has been reported to be highly conserved in class I proteins [36]. Class I catalytic domain is characteristic of a five strand parallel sheets flanked by α-helices with amino acid and ATP binding sites on opposite sides of a pseudo-2-fold symmetry. The Rossmann fold, in all Class I proteins has a connective polypeptide I (CPI) insert which is characterized by alpha and beta folds [40]. The conserved Motifs 1 and 2 across the class are present in the catalytic domains [40]. Detailed analysis of each protein family showed conserved motifs specific to each family (Additional file 3). Further, some conserved motifs unique only in the plasmodium proteins were observed (Figure 4 & Additional file 3 & 4).

**Figure 5:**
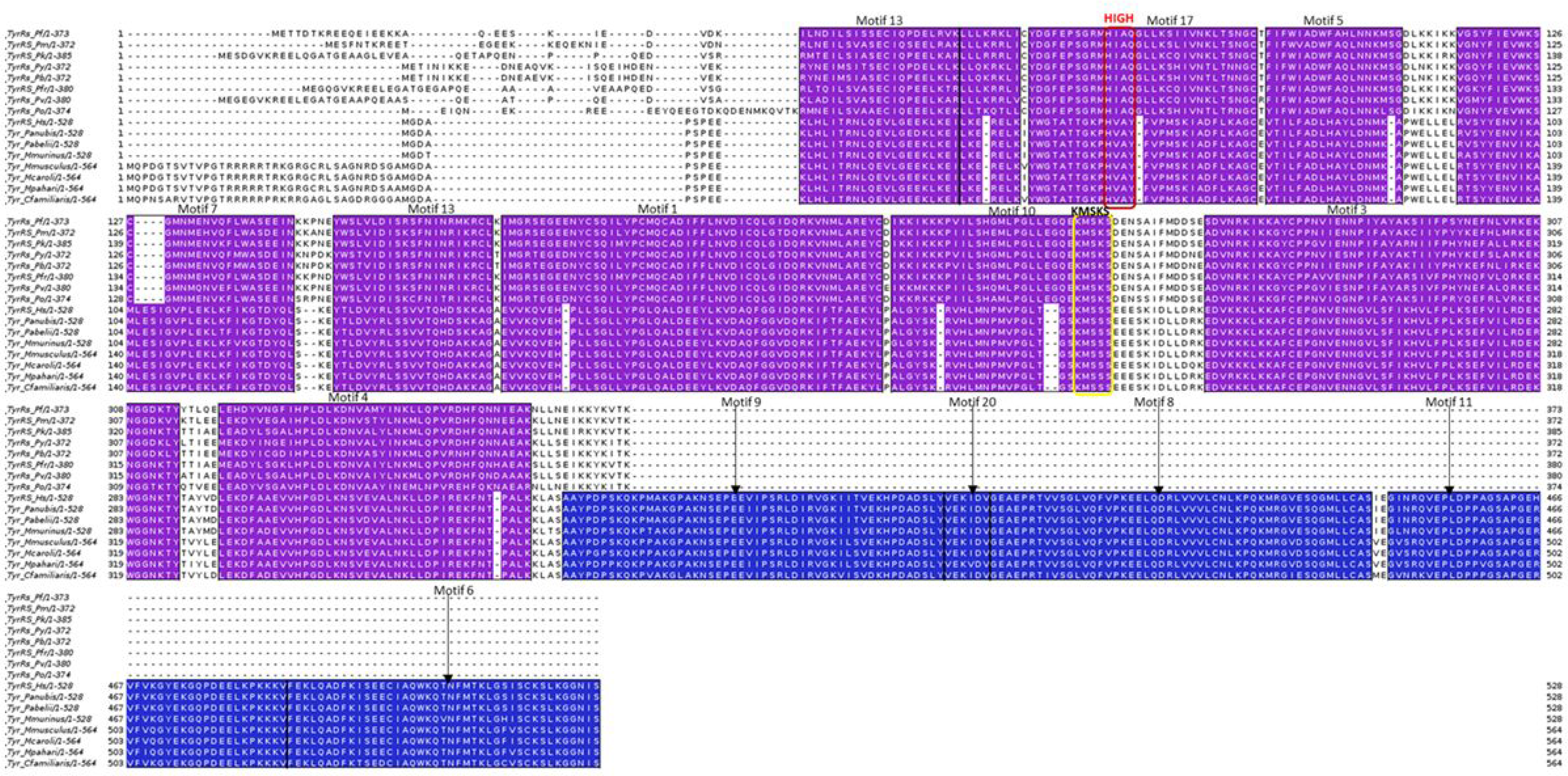
Motifs discovered in TyrRS family mapped to the multiple sequence alignment results. Motif numbering is based on MEME results. A purple colour shows motifs conserved in all sequences while motifs only present in mammalian sequences are shown in blue. The highly conserved HIGH and KMSKS motifs in Class II aaRSs are shown in a red and yellow box respectively.

**Figure 6:**
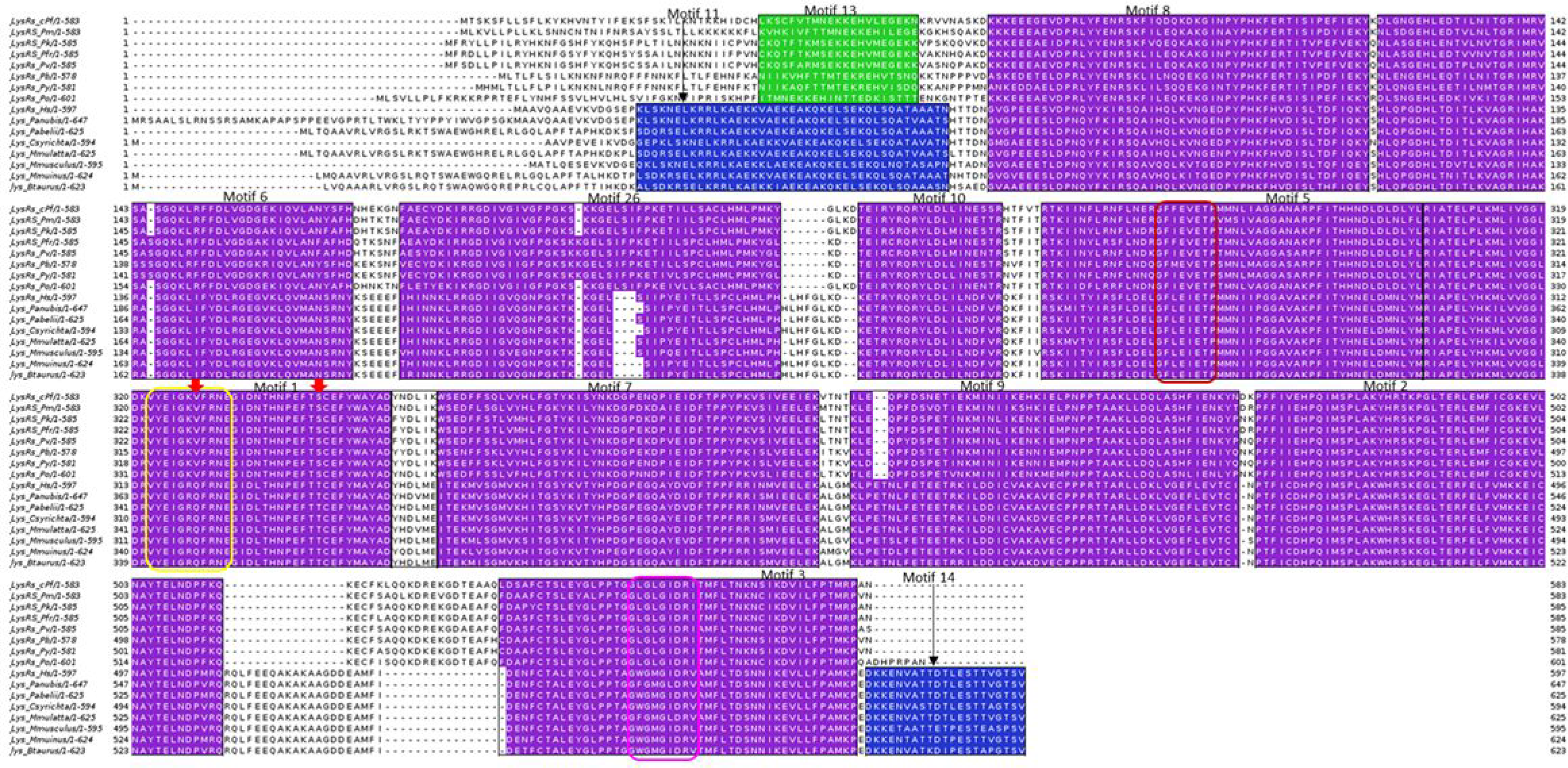
Mapping of discovered motifs in LysRS family to multiple sequence alignment. A purple colour shows motifs conserved in all sequences while motifs only present in mammalian sequences are shown in blue. One motif conserved only in plasmodium species is shown in green. Motif numbering is based on MEME results. The three conserved signatures in Class II aaRSs are shown in red, yellow and pink boxes. The red arrows show residues Val328 and Ser344 in *P. falciparum* which are key residues in binding of ATP.

On mapping the motifs to the multiple sequence alignments, differences at the residue level were observed despite the high level of motif conservation thus these residues can be the basis of drug discovery. Eukaryote specific motifs in ArgRS, MetRS, GluRS WHEP domain and AspRS are important for the association of proteins to a multi-tRNA synthetase complex in eukaryotes [112–114]. In human, nine aaRSs form a complex together with non-synthetase p18, p38 and p43 accessory proteins [114–116]. Leucyl, isoleucyl, glutaminyl, lysyl, methionyl, aspartyl, prolyl and glutamyl-tRNA synthetases form the multi-synthetase complex together with the auxiliary proteins in human aaRS but this complex is not present in plasmodium aaRSs [34].

These unique motifs may also play important roles other than the canonical catalytic roles [117]. Human LeuRS and GluRS, for example, have been reported to trigger leucine dependent cellular proliferation and glutamine dependent apoptosis by functioning as amino acid binding sensors [118,119]. Highly conserved motifs specific to each aaRS group are as a result of idiosyncratic insertions at the C-terminal or within or after the Rossmann fold of each protein family in this class [21,40,114] (Additional file 3 & 4). Methionine, valine, isoleucine and leucine aaRSs are all known to be specific to substrates that have aliphatic side chains and Motifs 20, 24 and 65 that are highly conserved in these four proteins may have a role in this specificity [40]. LeuRS, IleRS, MetRS, ArgRS, ValRS and CysRS have a structurally conserved anticodon binding domain characterized by α-helices and this may explain the conservation of Motifs 2, 20, 44 and 65 among these proteins (Figure 3) [40]. Plasmodium TrpRS has an N-terminal extension which is 227 amino acid residues long that constitute a AlaX-like domain and a linker region that function in binding of tRNA and in aminoacylation activity [120]. This extension is not present in the human TrpRS and thus explains the unique motifs at the N- terminal of the plasmodium proteins. Plasmodium sequences also have a lysine-enriched insertion at the C-terminal end of the KMSKS motif which is 15 residues long in PfTrpRS which is absent in the human sequence [120]. The domain for binding anticodons in Class I is located at the carboxyl terminal except for LeuRS. The structures of this region are highly divergent even within the sub-classes and is known to play an important role in tRNA discrimination [40].

In Class II, there were three highly conserved motifs across the class (Figure 4, Table 2). In the reporting of motif results of this class motif names are based on MEME results and not on previous literature. Motif 1 was present in 60 sequences, Motif 2 was present in 58 sequences while motif 20 was present in 76 sequences out of 89 sequences (Figure 4). Motif 1, motif 2 and Motif 19 discovered in Class II identified in this study contain the conserved signatures of Class II proteins (motif III, motif II and motif I respectively) reported by Chaliotis et al (2016) [36]. In Class II, motifs also clustered based on the protein family. Motif conservation among proteins may mean that these regions play a specific function in the proteins. Motif discovery was then done for each protein to determine conserved motifs within homologous sequences of each protein and the results presented as heat maps (Additional file 3).

**Table 2.**
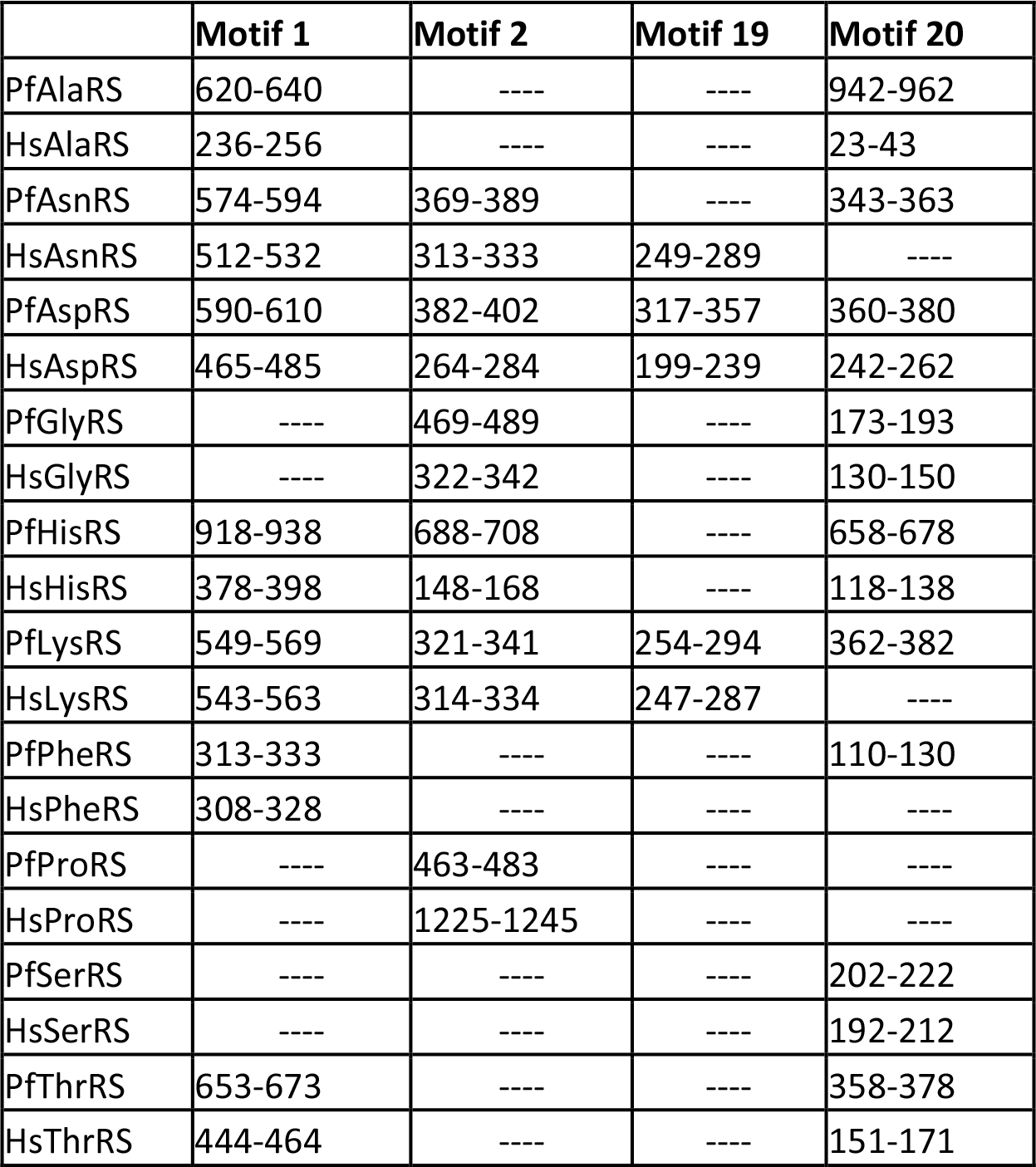
Highly conserved motifs in Class II aaRS. Motif 1, 2, 19 and 20 in Class II *P. falciparum* aaRS and the human homologues. Motif positions in the sequences are indicated and dashes are used where the motif is not present.

Class II aaRS have a highly conserved catalytic domain that occurs as β-sheet strands with α-helices on either side. This domain binds ATP, amino acid and the tRNA during aminoacylation. Motif 1 has been reported previously (as motif III) to be part of the active site forming α-helices and β-strands [36,121]. Motif 2, (Figure 4) also found at the catalytic site of proteins in this group forms β strands in pairs joined by a loop [35]. Motif I plays a role in binding of ATP while Motif 2 couples ATP, tRNA and amino acid binding [35,122]. Another weakly conserved motif in the active site of these proteins forms an α-helix that is linked to a β-strand with a proline residue at the end (Motif 19, Figure 4). This motif is known to be crucial in formation of dimers in most proteins of this class [123].

Further, subclasses in this class have conserved motifs within each subclass (Figure 4). For example, Ser, Thr, Gly, Pro and His aaRSs all belong to the Class IIa and have anticodon binding domains that are specific to the subclass [48]. These proteins are specific to small and hydrophobic amino acids and have motifs that are conserved among them as shown in the heatmap (Figure 4). The anticodon binding domain comprises of three α-helices five and β-stranded sheets and occurs in the C-terminus of this sub-class [48,124,125]. The anticodon binding domain is absent in SerRS as this protein does not require an anticodon to discriminate its cognate tRNA [47,48]. Subclass IIb which comprises of AsnRS, LysRS and AspRS have a unique anticodon binding domain at the N-terminal and share conserved motifs (Figure 4) [50,126,127]. This subclass of enzymes is specific to large polar and charged amino acid substrates and are similar in structural organization. AspRS is capable of catalysing aminoacylation of aspartate and asparagine and thus it can be classified as discriminating and non-discriminating protein just like GluRS [128,129]. Non-discriminating AspRS is only present in bacteria and archaea but not in eukaryotes [130]. Family specific motifs, can be attributed to the diversity in accessory domains found at the N- and C-terminal or within loops in the core domain [131].

#### Multiple sequence alignment and Motif mapping

Plasmodium and mammalian sequences for every aaRS family were aligned using TCOFFEE as indicated in the methodology. The alignment results were visualized using Jalview software and motifs discovered for each family mapped to these alignments [100]. A purple colour was used for the motifs that were conserved in all plasmodium and mammalian sequences, blue colour for only motifs conserved in mammalian species and green colour for motifs conserved only in plasmodium sequences (Additional file 4). On carrying out motif analysis and sequence alignment of Class I aaRSs, it was observed that not all families had the KMSKS signature though all proteins had the HIGH signature (Additional file 4). Alignment of ArgRS showed inserts in mammalian ArgRS at both the C- and N-termini that are not present in plasmodium sequences (Additional file 4A). The highly conserved HIGH signature in Class I aaRSs catalytic domain was observed in Motif 1 of this family (HVGH) (Additional file 4A). Motifs 10 and 12 which were conserved only in mammalian sequences were observed in the N-terminal. Human ArgRS has a basic 72 residue extension at the N-terminal which is characteristic of mammalian ArgRS and plays a role in interaction with accessory proteins like p43 to form the multi-synthetase complex [116,132]. Mammals also have an ArgRS isoform that lacks this extension and is believed to be important in ubiquitin dependent protein degradation where it forms Arg-tRNA^Arg^ which is transferred to ArgRS which then adds the arginine to all acidic N-terminal amino acids [133,134].

CysRS sequence alignment and motif mapping showed a highly conserved core domain and weakly conserved N- and C-terminal domains. The highly conserved HIGH signature was found in Motif 2 of this family occurring as HLGH in plasmodium and HMGH in the mammalian sequences (Additional file 4B). Motif 8, 10, 12, 13, 18 and 19 were conserved only in mammalian sequences while Motif 11 and 15 were only conserved in plasmodium sequences analysed in this family (Additional file 4B). GlnRS alignment also showed low conservation on both termini with inserts observed for the mammalian sequences at the N-terminal (Additional file 4C). Only two plasmodium specific motifs were found at the core domain, Motif 23 at the N-terminal end of the highly conserved HIGH signature (Motif 2) and Motif 29. Motif 8, 9, 11 and 13 were found only in the mammalian species (Additional file 4C). *P. falciparum* is reported to have Glutathione-S-transferase (GST)-like domains though their function in the malarial parasite has not been reported [34]. These domains are important in formation of multi-synthetase complex through protein-protein interactions in eukaryotes [21,22,117]. GST-like domains have also been reported in MetRS though just like in GlnRS, the function of these domains in plasmodium is not known unlike in eukaryotes where they play a role in protein-protein interactions [19].

The GluRS family also showed low conservation at the N-terminal with Motif 16 present in mammalian sequences at this terminal (Additional file 4D). The HIGH signature was found in Motif 3 as HIGH in all sequences analysed except for PfGluRS where it was HVGH (Additional file 4). *P. falciparum* GluRS sequence has a glutamine rich N-terminal from residue 68 as opposed to other plasmodium species. In mammals including human, this enzyme is a bifunctional protein acting both as GluRS and ProRS thus it catalyses aminoacylation of both proline and glutamate [135]. On alignment with plasmodium GluRS, the mammalian sequences showed a C-terminal extension indicating that it is the N-terminal end that catalyses glutamate aminoacylation. The human enzyme contains three motifs that link the two catalytic domains that function in formation of the multicomplex synthetase and play a role in protein-nucleic acid interactions [135,136]. Similar motifs have been reported in other aaRS like GlyRS, HisRS and TrpRS though they occur at the N-or C-termini of the core domains as a single copy as opposed to the Glu/ProRS where they occur as tandem repeats linking the two catalytic domains [135–137]. Human IleRS has an extension at the C-terminal which was absent in plasmodium sequences, but the core domain of this family was highly conserved (Additional file 4E). Motif 19, 20 and 26 were conserved in the C-terminal of mammalian sequences but absent in plasmodium sequences. The three tandem motifs in the human bifunctional Glu/ProRS have been shown to interact with two repeated motifs in IleRS at the C-terminal extension [138]. In IleRS, the HIGH signature was found in Motif 1 while the KMSKS signature was in Motif 3 occurring as HYGH and KMSKR respectively (Additional file 4E). Alignment and motif discovery of LeuRS family showed that this family of protein has low conservation even at the core domain (Additional file 4F). Motif 21, 25 and 27 were conserved in plasmodium sequences. Only Motifs 3, 5, 6, 26 and 36 were conserved through all mammalian and plasmodium sequences (Additional file 4F). The other motifs were conserved only in mammalian sequences. The highly conserved Motif 6 had the HIGH signature occurring as HVGH for PfLeuRS, PmLeuRS, PoLeuRS, PyLeuRS, HMGH for PfrLeuRS, PvLeuRS and PkLeuRS and HLGH in the analysed mammalian sequences (Additional file 4F). Anticodon binding domain in LeuRS is located at the C-terminal which had a low conservation as seen in Additional file 4F and this may provide specific targets for drug discovery [139]. Motif discovery and alignment of MetRS showed high conservation of mammalian sequences. Some unique motifs were only present in plasmodium MetRS but were absent in mammalian sequences (Additional file 4G). The highly conserved HIGH signature was observed in Motif 8 which was conserved in all sequences analysed while the KMSKS signature was found in Motif 14 conserved in plasmodium and Motif 6 in mammalian sequences (Additional file 4G). The catalytic domain of MetRS was weakly conserved with only Motif 1, 2, 4, 8, and 15 being conserved in all sequences at this region. The C-terminal showed mammalian and plasmodium specific motifs. Motif 5 and 9 found at the N-terminal were conserved in all analysed sequences in this family (Additional file 4G).

TrpRS alignment revealed a plasmodium specific extension at the N-terminal characterised by Motif 8, 9, 10 and 14 (Additional file 4H). This extension plays a role in aminoacylation and tRNA binding as reported in *P. falciparum* [34]. In *P. falciparum*, this extension comprises of a linker region and an AlaX-like domain that plays a role in tRNA binding but does not edit mis-acylations as observed with *Pyrococcus horikoshii* [120]. The core domain and the C-terminal of TrpRS family showed highly conserved motifs in all the sequences with only a short Motif 18 present in mammalian sequences (Additional file 4H). Alignment and mapping motifs discovered in TyrRS sequences showed high conservation of motifs at the core domain (Figure 5). Alignment of sequences in this family showed an extension at the C-terminal of the mammalian TyrRS which was missing in all plasmodium sequences (Figure 5). This extension was characterised by Motifs 6, 8, 9, 11 and 20 which were conserved in all the mammalian sequences analysed (Figure 5). This extension in human TyrRS is an endothelial monocyte-activating polypeptide II (EMAPII) domain that has cytokine-like functions like angiogenesis and inflammation [22,140]. Motif discovery showed that the core domain is highly conserved across the mammalian and plasmodium TyrRS sequences (Figure 5). The catalytic domain of the human sequence is also different from the malarial parasites in that it has a buried tripeptide cytokine motif (Glu-Leu-Arg) while in plasmodium this motif is on the surface [22,53]. ValRS alignment showed a N-terminal extension for the mammalian sequences that was absent in all plasmodium sequences comprising of Motifs 14, 16, 18, 22 and 25 (Additional file 4J). Mapping of motifs showed that the catalytic domain of proteins analysed in this family are highly conserved though a few plasmodium specific motifs were observed. The highly conserved HIGH signature was found in Motif 2 of this family while the KMSKS signature was in Motif 7 (Additional file 4J). The N-terminal domain showed Motif 16, 33, 34 and 35 which were conserved only in plasmodium sequences (Additional file 4J). Motifs 20, 30 and 38 that were specific to mammalian sequences were also observed at the N-terminal (Additional file 4J).

Alignment of AlaRS sequences showed a N-terminal extension of varying lengths in the plasmodium species which was absent in mammalian AlaRS (Additional file 4K). The C-terminal of the proteins in this family showed Motifs 20, 21 and 29 that were only conserved in plasmodium sequences and not in human as well as mammalian specific motifs (Motif 8, 14, 17 and 18). AsnRS, LysRS, and AspRS alignment and motif discovery showed low conservation at the N-terminal while core domains and the C-terminal showed high conservation. The anticodon binding domain of these proteins is located at the highly variable N-terminal and thus drugs that specifically bind to the parasite tRNA binding site can be designed [14,141]. Motif 11, 12 and 17 were conserved in plasmodium sequences of AsnRS family at the N-terminal while in this region, Motif 5, 6 and 13 conserved in mammalian sequences were observed (Additional file 4L). In AspRS both the catalytic domain and the C-terminal were highly conserved with the presence of two short Motifs (16 and 20) conserved only in mammals (Additional file 4M). GlyRS, HisRS, ProRS, ThrRS families belong to the subclass IIa and have a highly conserved tRNA binding region at the C-terminal as seen in the alignments and motifs in this region (Additional file 4 N, O, R and T). HisRS family showed a N-terminal extension for all plasmodium sequences analysed but absent in the mammalian sequences (Additional file 4O). This extension was characterised by Motifs 11, 12, 14, 15, 17, 18, 19 and 23 (Additional file 4O). However, SerRS which also belongs to this subclass does not need an anticodon to discriminate its substrate and thus lacks this domain [48] and the C-terminal of this family showed low conservation (Additional file 4S). ProRS showed Motif 17 and 20 which were conserved only in plasmodium sequences analysed (Additional file 4R). Plasmodium ProRS has a Ybak domain at the N-terminal which edits mischarged Pro-tRNA^Ala^ and Pro-tRNA^Ser^ and this may explain the plasmodium specific motifs at the N-terminal [15,34,57]. The mammalian sequences analysed for this family were of the cytosolic bifunctional Glu/ProRS proteins and this explains the mammalian specific motifs observed at the N-terminal which is believed to be the region responsible for glutamate aminoacylation (Additional file 4R).

PheRS motif discovery and alignment showed that the plasmodium sequences are highly variable when compared to mammalian PheRS (Additional file 4Q). Motifs 9, 10, 11 and 13 were conserved only in plasmodium while Motifs 5, 6, 8 and 14 were conserved in mammalian sequences in this family (Additional file 4Q). Only Motifs 1, 2, 3 and 4 were conserved across all the sequences in this family (Additional file 4Q). Plasmodium PheRS has a nuclear localization signal and DNA binding domains and thus in addition to aminoacylation, this enzyme mediates cellular processes by binding DNA [142]. Despite high conservation at the aaRS active sites, differences were noted at the residue level after the sequences were aligned. For example, in LysRS family, *P. falciparum* ATP binding pocket at positions Val328 and Ser344 corresponds to Gln321 and Thr338 respectively in the human protein (Figure 6). Residues with a large side chain at this position like observed in human LysRS do not favour binding of cladosporin a known inhibitor for PfLysRS [25]. These two residues are thus believed to be responsible for selective binding of cladosporin to *P. falciparum* and not human LysRS [25]. Discovery of drugs that have high specificity to parasitic proteins has for a long time been a challenge resulting in drug toxicity in human cells [14]. The alignment results showed striking differences at the sequence level of plasmodium and human aaRSs that can further be explored for the design and development of drugs with few side effects.

#### Phylogenetic tree calculations and pairwise sequence identity calculations agree in grouping sequences

On conducting phylogenetic tree analysis, all plasmodium species clustered together, and this was also seen on performing all versus all pairwise sequence identity calculations (Figure 7, Figure 8 and Additional file 6). In this study, numbering of sequences in sequence identity heatmaps was based on the branching of phylogenetic trees. In Class I, plasmodium sequences in TyrRS family showed the highest sequence identity (above 85%) while GlnRS plasmodium sequences showed the lowest sequence (below 75%) identity among plasmodium families. In most of the families, *P. yoelii* and *P. bergei* sequences were clustered together in the trees. *P. vivax*, *P. fragile* and *P. knowlesi* were also clustered together in many families, indicating that they are highly conserved and share evolutionary history. These similarities were also captured in sequence identity calculations, and reflected as imaginary boxes in heat maps. Here we will name them as “conservation boxes”. *P. bergei* and *P. yoelii* are rodent malaria parasites and are used to study human malaria [143,144]. *P. fragile* infects simians and studies have shown that human red blood cells do not support the growth of this parasite but it showed a high sequence identity to *P. knowlesi* whose natural vertebrate host is *Macaca fascicularis* but has been reported to infect human in some parts of Southeast Asia [145,146]. *P. knowlesi* has been reported to have a close phylogenetic relationship to *P. vivax* [147] and the two showed a sequence identity above 95% in TyrRS (Figure 8). *P. fragile*-monkey models can thus be used to study parasite-host-system for the immunological response of the falciparum-like parasite both *in vivo* and *in vitro* [148].

**Figure 7.**
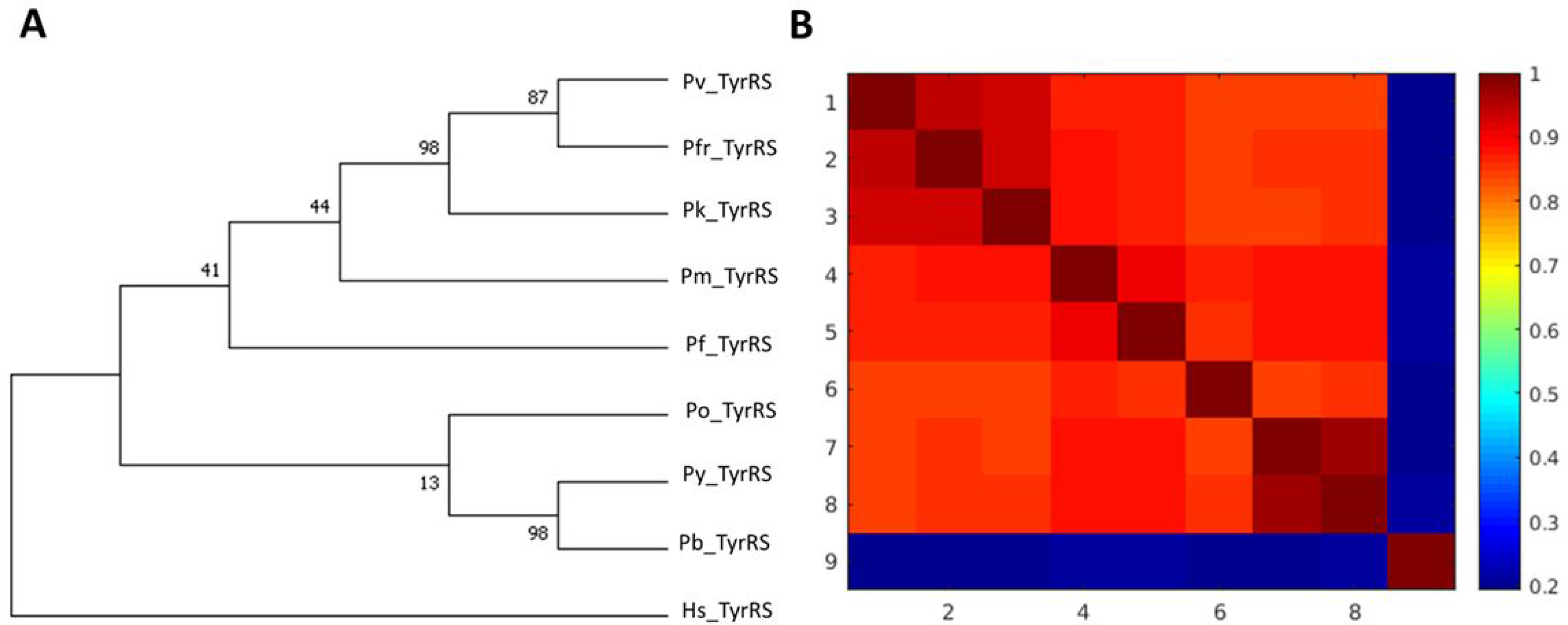
**A)** TyrRS family phylogenetic tree. Maximum Likelihood method was used to infer evolutionary history using Le_Gascuel_2008 model at 95% site coverage [149]. Phylogenetic tree calculations were done using MEGA7 [101]. The tree that had the highest log likelihood (−2978.09) is shown. Initial tree(s) for the heuristic search were obtained by using BioNJ and Neighbor-Join algorithms to a matrix of pairwise distances calculated using a JTT model, and then selecting the topology with higher log likelihood value. A Gamma distribution was used to calculate evolutionary rate differences among sites (5 categories (+*G*, parameter = 0.4355)). Nine amino acid sequences were used for this analysis. There were 343 positions after the calculations. **B)** TyrRS pairwise sequence calculations. The sequence identity values of the sequences in the TyrRS family is shown. The heatmap shows the identity scores as a color-coded matrix for every aaRS sequence versus every aaRS sequence in this family. Conservation increases from blue to red in the heat map.

**Figure 8.**
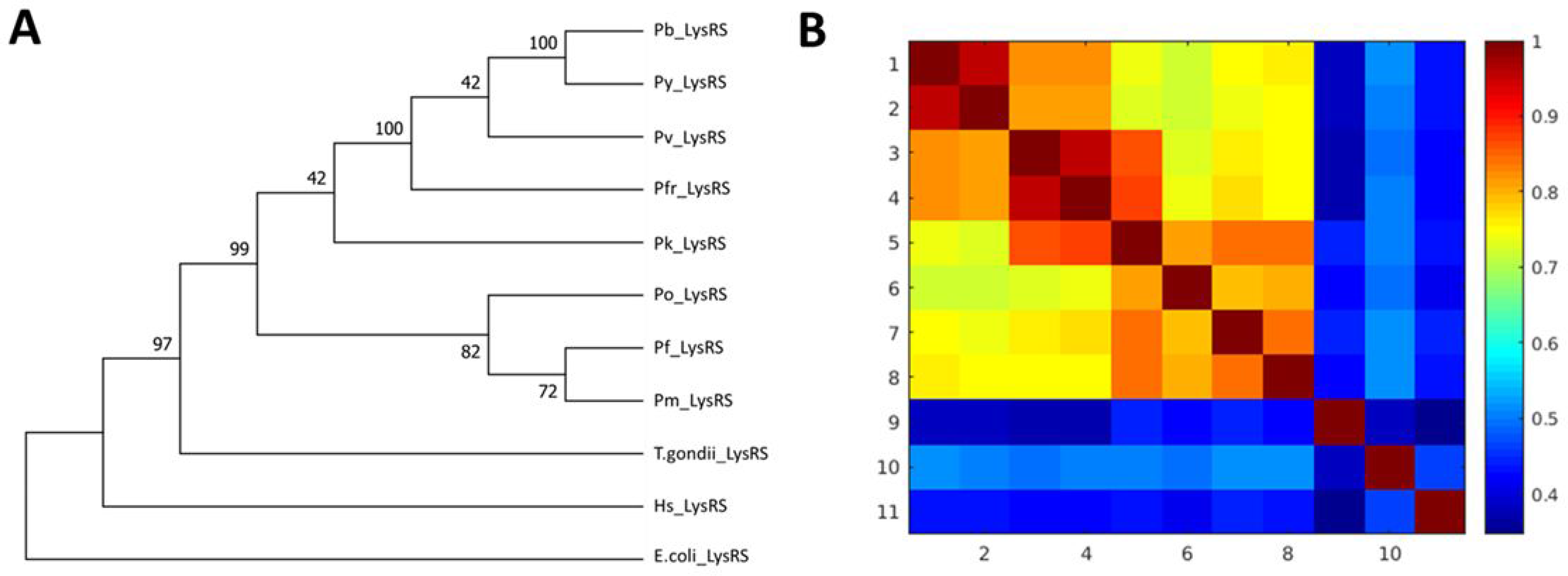
**A)** LysRS family phylogenetic tree. Maximum Likelihood method was used to infer evolutionary history using Le_Gascuel_2008 model at 90% site coverage [149]. Phylogenetic tree calculations were done using MEGA7 [101]. The tree that had the highest log likelihood (−6116.25) is shown. Initial tree(s) for the heuristic search were obtained by using BioNJ and Neighbor-Join algorithms to a matrix of pairwise distances calculated using a JTT model, and then selecting the topology with higher log likelihood value. A Gamma distribution was used to calculate evolutionary rate differences among sites (5 categories (+*G*, parameter = 0.6075)). Eleven amino acid sequences were used for this analysis. There were 503 positions after the calculations. **B)** LysRS pairwise sequence calculations. The sequence identity values of the sequences in the LysRS family is shown. The heatmap shows the identity scores as a color-coded matrix for every aaRS sequence versus every aaRS sequence in this family. Conservation increases from blue to red in the heat map.

In ArgRS sequence identity calculations, plasmodium sequences had above 80% sequence identity and motif discovery showed that all motifs identified were conserved in all sequences (Additional file 3 & 6.1). ValRS plasmodium sequences showed 80% sequence identity with PvValRS, PkValRS and PfrValRS clustering together with a 90% sequence identity. In this family, PyValRS and PbValRS showed above 95% sequence identity, clustered together in the phylogenetic tree and shared Motif 36 which was absent in the other plasmodium sequences (Additional file 3 & 6.11). PvCysRS, PkCysRS and PfrCysRS clustered together with a 90% sequence identity and shared Motif 22 which was missing in other plasmodium sequences (Additional file 3 & 6.2). Motif 27 was present only in PyCysRS and PbCysRS and these two sequences showed a 90% sequence identity Additional file 3 & 6.2). PfrGlnRS, PkGlnRS and PvGlnRS clustered together and Motif 34, 35 and 37 were only present in these sequences (Additional file 3 & 6.3). In this family, *E. coli*, human and *S. cerevisiae* sequences formed an outgroup showing they are the oldest aaRS (Additional file 6.3). GluRS plasmodium sequences had above 75% sequence identity and shared all identified motifs (Additional file 3 & 6.4). PbIleRS and PyIleRS shared Motif 38 and 39 and showed 95% sequence identity (Additional file 3 & 6.5). Cryptosporidium and toxoplasma belong to the apicomplexan family together with plasmodium and their sequences showed about 50% sequence identity to plasmodium sequences in IleRS and MetRS family (Additional file 6.5 & 6.7). PvLeuRS, PfrLeuRS and PkLeuRS had 80% sequence identity and shared Motif 39 (Additional file 3 & 6.6). In TrpRS, Motif 21 and 23 were only identified in PbTrpRS and PyTrpRS which had 90% sequence identity. In all families in Class I, human sequences showed low sequence identities (below 40%) compared to the plasmodium sequences (Additional file 6).

In Class II aaRSs, ProRS family showed the highest sequence identity with plasmodium sequences having above 80% sequence identity (Additional file 6). The high sequence identity among plasmodium sequences was also reflected in motif identification where all the sequences shared the identified motifs (Additional file 3). In Class IIa GlyRS showed the least conservation with most of the sequences having less than 65% sequence identity (Additional file 6.16). GlyRS family showed low conservation with sequence identity less than 70% for all sequences except for PfrGlyRS, PvGlyRS and PkGlyRS which formed a conservation box with a sequence identity of about 75% (Additional file 6.16). This clustering was also seen in motif identification whereby PfrGlyRS, PvGlyRS and PkGlyRS had Motif 24 which was absent in all other plasmodium sequences in this family. PbGlyRS and PyGlyRS had a sequence identity of 90% and shared Motif 27, 30 and 34 (Additional file 3 & 6.16). *P. falciparum* ThrRS had a low sequence identity compared to other plasmodium sequences and it also branched separately in the phylogenetic tree. In SerRS family, human, *T. brucei*, *C. albicans*, *T. gondii* and *C. parvum* also formed a conservation box but with a sequence identity of about 65%. *P. vivax*, *P. fragile* and *P. knowlesi* in this family had a high sequence identity forming a conservation box and clustered together in the phylogenetic tree (Additional file 6.20). PfrThrRS, PvThrRS and PkThrRS shared Motif 24, 27, 29 and 33 showing these sequences are closely related as depicted by trees and sequence identity calculations (Additional file 3 & 6.20). In SerRS family, plasmodium sequences formed a conservation box with about 75% sequence identity with each other except for *P. yoelii* which was more identical to *P. bergei* with a sequence identity of 90% (Additional file 6.19). In motif identification, *P. yoelii* shared Motif 20 and 22 which were all absent in all other plasmodium sequences explaining the high sequence conservation (Additional file 3). PkSerRS and PvSerRS branched together and the two shared Motif 19 showing that the sequences are closely related (Additional file 3). In HisRS family, all plasmodium sequences formed a conservation box showing more than 70% sequence identity to each other except for PfHisRS (Additional file 6.14). This difference was also seen in motifs identified in this family where Motif 21 was present in all plasmodium sequences but absent in PfHisRS (Additional file 3).

In Class IIb, AsnRS sequences were highly conserved with above 80% sequence identity while AspRS was the least conserved with about 65% sequence identity (Additional file 6.13 & 6.15). The high sequence conservation in AsnRS was also seen in motif discovery where all plasmodium sequences shared identified motifs (Additional file 3). In AsnRS family, *C. ubiquitum* showed a higher sequence identity to *S. typhi* sequence than to *T. gondii* which belongs to the same phylum. PvAspRS and PkAspRS branched together in tree calculation and these two proteins shared Motif 22, 26 and 28 showing they are closely related (Additional file 3 & 6.15). In LysRS family, plasmodium sequences showed a sequence identity of above 75% with PbLysRS and PyLysRS forming a conservation box with about 95% sequence identity (Figure 8). PfrLysRS, PkLysRS and PvLysRS also formed a conservation box and these three proteins shared Motif 15 which was absent in other plasmodium sequences (Figure 8, Additional file 3).

Overall, PheRS was the least conserved family in Class II with plasmodium sequences with only about 50% sequence identity and this was seen during motif discovery where only a few motifs were conserved across species (Additional file 3 & 6). In the AlaRS family, *P. falciparum* (sequence 5 in the heatmap) was less conserved compared to other plasmodium sequences as seen in the (Additional file 6). Plasmodium sequences in AlaRS family showed a sequence identity above 70% (Additional file 6.12). In this family, *P. vivax*, *P. fragile* and *P. knowlesi* also formed a conservation box while *P. yoelii* and *P. bergei* also formed a conservation box indicating that these sequences are highly conserved compared to other plasmodium sequences. PfrAlaRS, PvAlaRS and PkAlaRS shared Motif 31 which was absent in all other plasmodium sequences but present in mammalian sequences (Additional file 3). In all the families in Class II, human sequences in this class branched out as an out group and this is supported by the low sequence identity (below 40%) shown in the conservation heatmaps (Additional file 6).

### PART 2 - STRUCTURAL ANALYSES

#### Accurate 3D protein models are calculated for Class I and Class II aaRSs

In the PDB, there are only four Class I (ArgRS, MetRS, TrpRS, TyrRS) and two Class II (LysRS and ProRS) structures that were available with reasonable quality. As a first step, each of these crystal structures was remodelled to eliminate the missing residues, except PfTyrRS, as this structure does not have missing residues. It was previously shown that homology modelling with a very high sequence template identity (or remodelling itself) does not introduce modelling errors [150]. As a next step, these models were used to model the 3D structures of the homologues (see Additional file 2 for further information).

For each protein, 100 homology models were calculated, and the three best models selected based on z-DOPE scores. DOPE score is an atomic statistical potential which depends on a native protein structure [151]. It is highly accurate in assessment of the quality of protein models as it accounts for the spherical and finite shape of the protein native structure [151–153]. It depends on the number of atom pairs considered and thus the number of all possible pairs of heavy atoms in the protein are normalized to get the z-DOPE score [151,154]. Models with lowest z-DOPE were selected and model quality assessment was done using Verify 3D [96], ProSA [95] and QMEAN [97] webservers. Verify 3D assesses the compatibility of the 3D structure with the amino acid sequence (1D) and assigns a class to the structure based on the local environment, location and secondary structure and compares this to known native structures [96]. At least 80% of the amino acid residues should have a score greater than or equal to 0.2 in the 3D/1D profile for the structure to be considered of good quality. ProSA-web is a tool for checking errors in a 3D model and displays the quality score as graphical presentation. Areas of the model that are not accurate are identified by a plot of local quality scores which are then mapped on the 3D structure using colour codes [95]. QMEAN score describes the major geometrical aspects of protein models using five structural descriptors. The overall status of residues is described by a solvation potential, long-range interactions are assessed by secondary structure-specific pairwise residue-level potential that is dependent on distance and a torsion angle potential is used to determine the local geometry which is calculated over three consecutive residues [97]. Descriptors of solvent accessibility and the agreement between calculated and predicted structures are also used in calculating the score [97]. All the calculated models passed the quality evaluation tests from these three tools (Additional file 2).

The models for the plasmodium ArgRS were built using 5JLD [89] as a template while 4ZAJ was used for the human homologue. The ArgRS models consist of the N-terminal, catalytic domain and the anticodon binding domain. All the models for MetRS, which included the catalytic domain and the anticodon binding domain, were calculated using 4DLP [90]. Plasmodium TrpRS models were built with 4J75 [155] while 1R6T [156] was used for HsTrpRS. It was possible to model the N-terminal, catalytic and anticodon binding domains for this family. The crystal structure 5USF [92] was used for the calculation of plasmodium TyrRS while 1Q11 [156] consisting only the catalytic and anticodon binding domain was used to model the HsTyrRS. The catalytic and anticodon binding domains of LysRS were built using 4DPG [94] as the starting structure while 4NCX [60] was used for building ProRS models which included a zinc-binding like domain at the C-terminal.

The 3D models were, then, used for mapping identified motifs to structures as well as for the search of alternate druggable sites in *P. falciparum* homologues.

#### Motif mapping to homology models

Out of all identified motifs (Additional file 3), the motifs of the six families with structures were mapped into the 3D structures (Figure 9, Figure 10 and Additional file 5). The start and end residues for motifs identified in the six families are shown for *P. falciparum* and the human homologues (Table 3). In ArgRS family, motifs were conserved in all analysed structures except Motif 16 which was present only in the plasmodium sequences but absent in in HsArgRS (Additional file 5A, Figure 9). HsArgRS N-terminal had Motifs 10 and 11 which were absent in plasmodium structures. Motif 13 was not positionally conserved in the analysed structures. In plasmodium it occurs in the anticodon binding domain and the N-terminal while in HsArgRS it occurs in catalytic and the anticodon binding domains (Additional file 5A). In HsMetRS, Motif 5 was in the anticodon binding domain while in plasmodium structures this motif was mapped to the catalytic site. The motif occurs in an alpha helix region in HsMetRS while in PfMetRS the site consists of beta sheets. Motif 14 occurring in the catalytic site and a loop region in PfMetRS was missing in HsMetRS structure (Figure 9). Motif 10 was present in HsMetRS anticodon binding domain but absent in plasmodium. Other motifs in this family were conserved across all analysed structures. In TrpRS, Motif 7 was only present in HsTrpRS but absent in all plasmodium structures. Motif 8, 9 and 10 were present only in PyTrpRS (Additional file 5C). Motif 1 and Motif 4 were mapped at the catalytic domain in all structures except in PyTrpRS where they are in the anticodon binding domain (Additional file 5C). Motif 2 was present at the catalytic domain of all the TrpRS homology model structures but absent in PyTrpRS. In TyrRS family, Motif 14 was conserved in PfTyrRS, PkTyrRS, PmTyrRS, PvTyrRS and PyTyrRS while Motif 12 was only present in human (Figure 9 & Additional file 5D).

**Table 3:** Starting and ending positions of motifs identified in PfArgRS, PfMetRS, PfTrpRS, PfTyrRS, PfLysRS and PfProRS as well as the human homologues. Dashes show where the motif was not present.

|  | Motif 1 | Motif 2 | Motif 3 | Motif 4 | Motif 5 | Motif 6 | Motif 7 | Motif 8 | Motif 9 | Motif 10 | Motif 11 | Motif 12 | Motif 13 | Motif 14 | Motif 15 | Motif 16 | Motif 17 |
| --- | --- | --- | --- | --- | --- | --- | --- | --- | --- | --- | --- | --- | --- | --- | --- | --- | --- |
| PfArgRS | 135-184 | 441-490 | 308-357 | 506-555 | 364-413 | 30-64 | 230-279 | 70-119 | 186-226 | ---- | ---- | ---- | 15-29 | 558-578 | 420-434 | 286-306 | ---- |
| HsArgRS | 198-247 | 499-548 | 371-420 | 568-617 | 428-477 | 101-135 | 293-342 | 142-191 | 249-289 | 50-99 | 620-660 | 8-48 | 346-360 | 478-498 | 549-563 | ---- | 363-370 |
| PfMetRS | 243-292 | 519-559 | ---- | 439-459 | 331-360 | ---- | ---- | 193-242 | 405-433 | ---- | ---- | 562-611 | ---- | 468--517 | 301-329 | ---- | 657-706 |
| HsMetRS | 285-334 | 598-638 | ---- | 510-530 | 711-740 | ---- | 460-509 | 234-283 | 35-63 | ---- | ---- | 104-153 | 159-208 | ---- | 343-371 | 839-888 | ---- |
| PfTrpRS | 301-350 | 447-496 | 512-561 | 249-298 | 392-441 | 562-611 | ---- | 74-123 | ---- | 124-173 | 1-41 | 203-243 | ---- | 43-71 | 354-368 | 612-626 | 174-199 |
| HsTrpRS | 154-203 | 304-353 | 354-403 | 102-151 | 247-296 | 404-453 | 48-97 | ---- | ---- | ---- | 222-242 | ---- | 6-46 | ---- | 206-220 | 454-468 | ---- |
| PfTyrRS | 172-221 | 60-88 | 265-314 | 320-360 | 91-108 | ---- | 116-144 | ---- | ---- | 223-251 | ---- | ---- | 30-50 | 149-169 | 254-264 | 362-469 | 51-58 |
| HsTyrRS | 150-199 | 39-67 | 240-289 | 295-335 | 69-86 | 488-528 | 97-125 | 395-444 | 339-388 | 200-228 | 447-487 | 1-29 | 129-149 | ---- | 229-239 | 89-96 | 30-37 |
| PfLysRS | 304-353 | 465-514 | 532-581 | 177-226 | 254-303 | 121-170 | 360-409 | 70-119 | 414-463 | 228-248 | ---- | 515-529 | 40-60 | ---- | ---- | 354-359 | ---- |
| HsLysRS | 297-346 | 459-508 | 526-575 | 169-218 | 247-296 | 114-163 | 353-402 | 63-112 | 409-458 | 221-241 | ---- | 509-523 | 577-597 | ---- | ---- | 347-352 | 1-15 |
| PfProRS | 362-411 | 268-317 | 662-711 | 502-551 | 570-606 | 447-496 | 319-359 | ---- | ---- | ---- | 718-746 | ---- | ---- | 418-446 | 615-648 | ---- | 109-149 |
| HsProRS | 1124-1173 | 1030-1079 | 332-381 | 1266-1315 | 1338-1374 | 1209-1258 | 1081-1121 | 447-496 | 666-715 | 392-441 | 1484-1512 | 191-240 | 257-306 | 1180-1208 | 1383-1416 | 1443-1483 | 497-537 |

**Figure 9:**
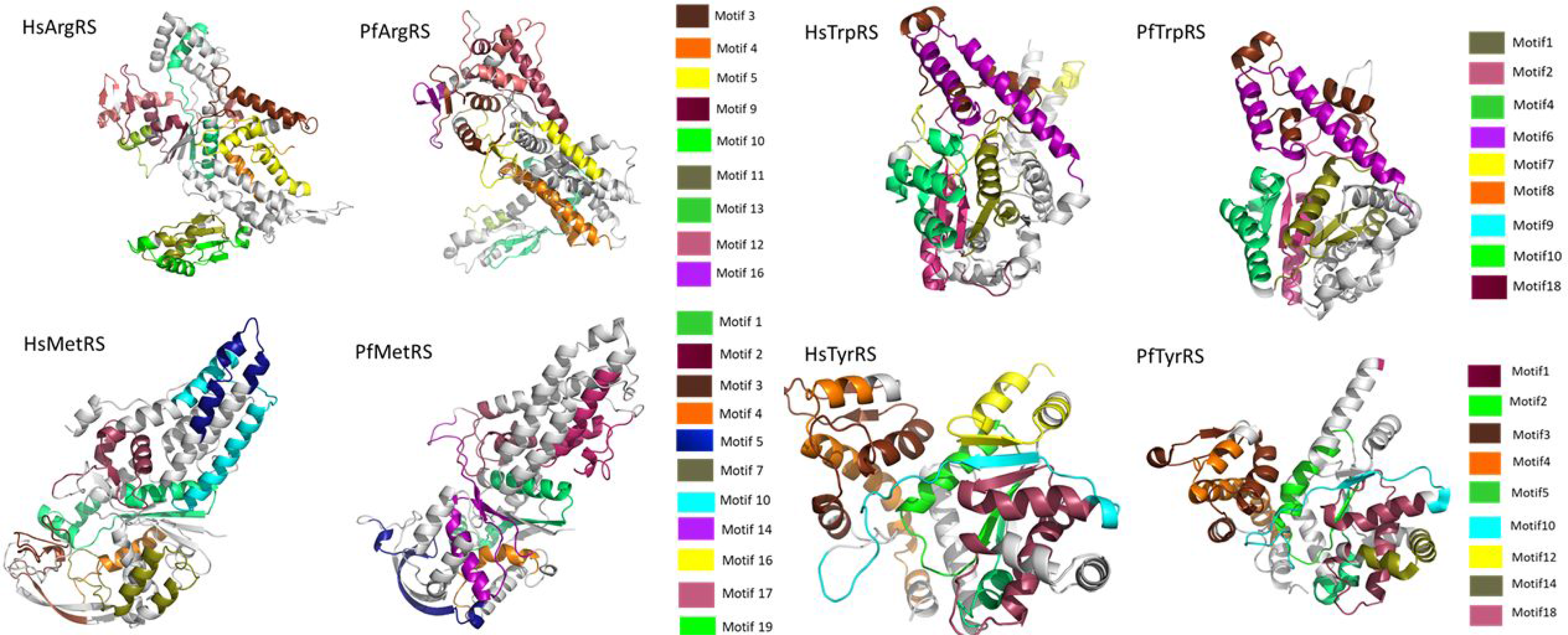
Class I motifs mapped to the homology models of *P. falciparum* and the human homologue. Motifs are numbered according to the MEME results.

**Figure 10:**
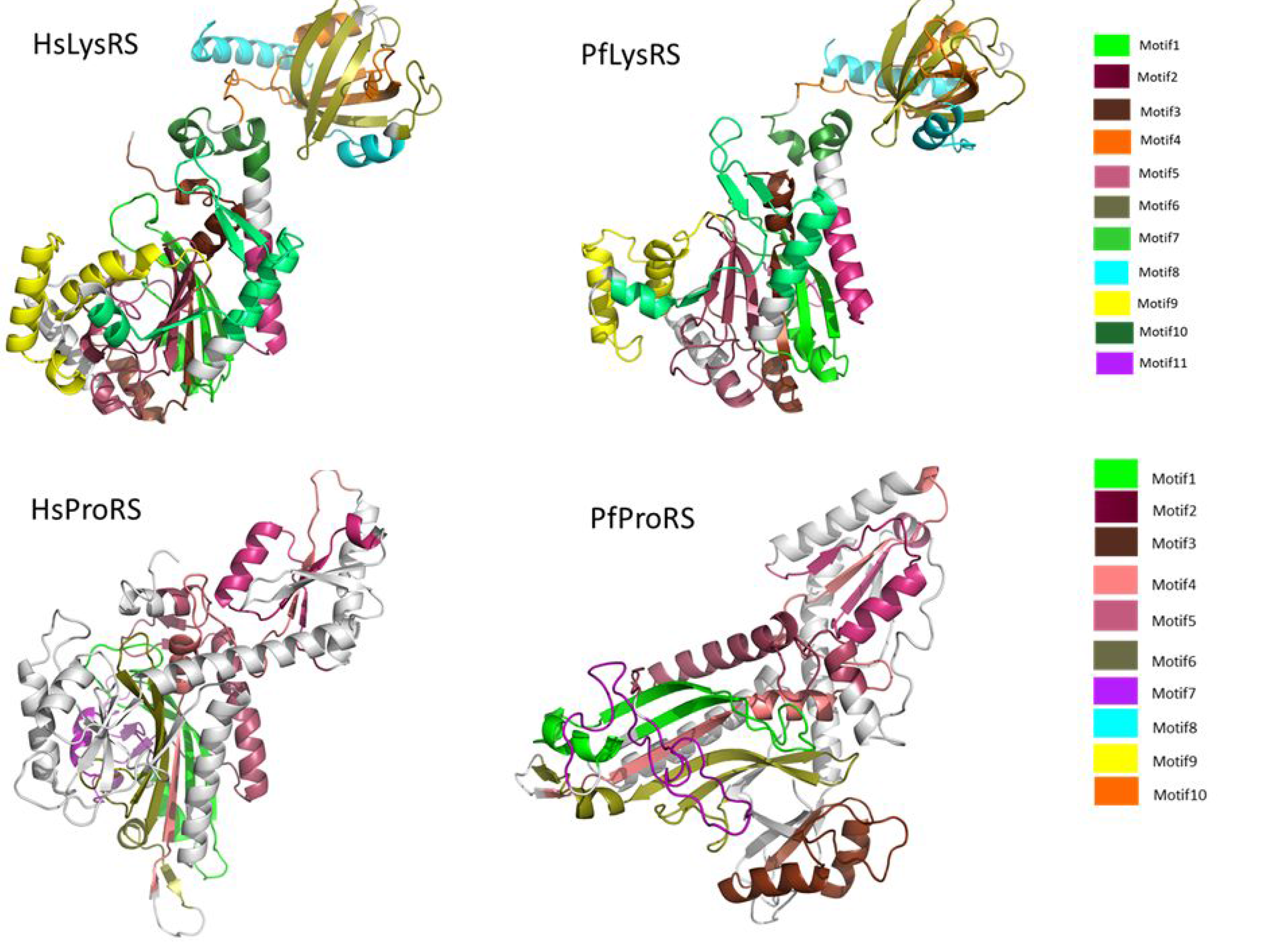
Class II motifs mapped to the homology models of *P. falciparum* and the human homologues. Motifs are numbered according to the MEME results.

In Class II, in LysRS, Motif 9 occurs at the catalytic domain in a region consisting of alpha helices and loops in all structures except PfrLysRS and PmLysRS where it mapped in a region consisting of both beta strands and alpha helices. PfrLysRS and PmLysRS did not have Motif 4 present in the anticodon binding domain of all other structures. Motif 8 mapped in a region consisting of alpha helices in all structures except in PfrLysRS and PmLysRS where the region consisted of beta sheets and alpha helices (Additional file 5E). Mapped motif in ProRS were conserved in all analysed secondary structures (Figure 10 & Additional file 5F).

#### New potential druggable sites in *P. falciparum* aaRSs are identified

FTMap provides information on binding hot spots and the druggability of these sites using probes from fragment libraries [107]. These fragment hits can be used in identification of hits from larger ligands. On the other hand, SiteMap predicts possible binding sites using an algorithm that assigns site points using geometric and energetic properties [105,106]. The site points are then grouped to give sites which are ranked based on a SiteScore computed based on size, hydrophobicity, exposure to the solvent and the ease of donating or accepting hydrogens. Both FTMap and SiteMap showed consistency in prediction of probable binding sites. In all the six modelled proteins, FTMap and SiteMap were able to predict the known active sites which consists of the ATP and amino acid binding sites as the highest ranked site (Figure 11 & 2). Alternative sites were also predicted in PfArgRS, PfMetRS, PfProRS, and HsProRS that can be targeted for design of new drug classes using both FTMap and SiteMap (Figure 11, 12 & 13). Since two tools show consistency in prediction of possible binding sites, we only discuss the results from FTMap in this study.

**Figure 11:**
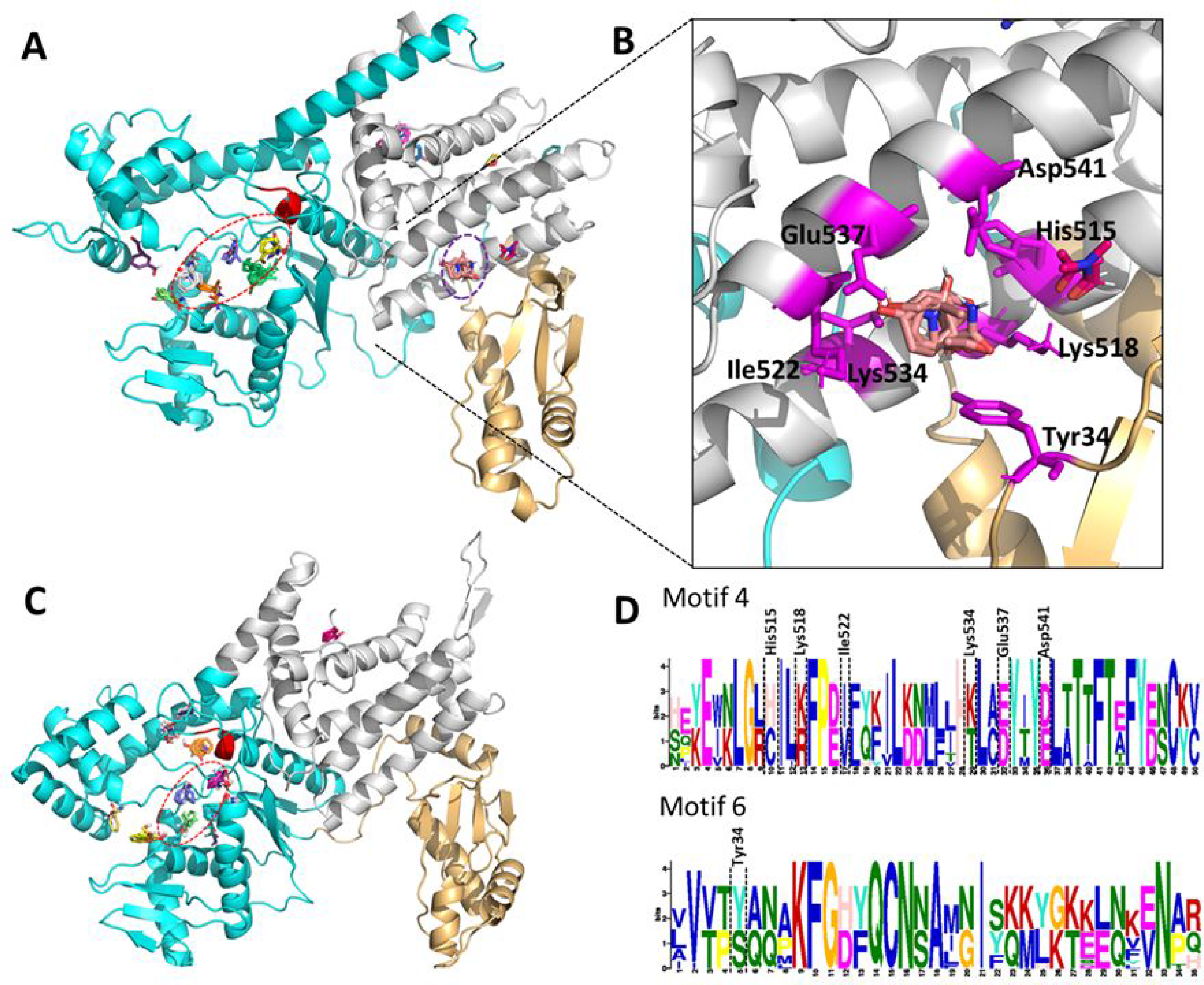
Homology models of PfArgRS and HsArgRS and prediction of potential ligand binding sites. The catalytic domain of the models is shown in cyan, the anticodon binding domain (ABD) in grey and the N-terminal domains of PfArgRS and HsArgRS are shown in a light orange colour. The HIGH and KMSKS motifs highly conserved in Class I are shown in red and yellow respectively. Known druggable sites are shown by the red dotted ellipses while the predicted site in PfArgRS by FTMap is shown in purple dotted ellipses. **A)** PfArgRS homology model. **B)** Insert - zoomed view of the predicted druggable site in PfArgRS with the residues interacting with probes represented as magenta sticks. **C)** HsArgRS homology model. No probable druggable sites were predicted in human homologue. **D)** Motif 4 and 6 logos showing conservation of residues in this family and PfArgRS residues interacting with probes at the predicted site.

**Figure 12:**
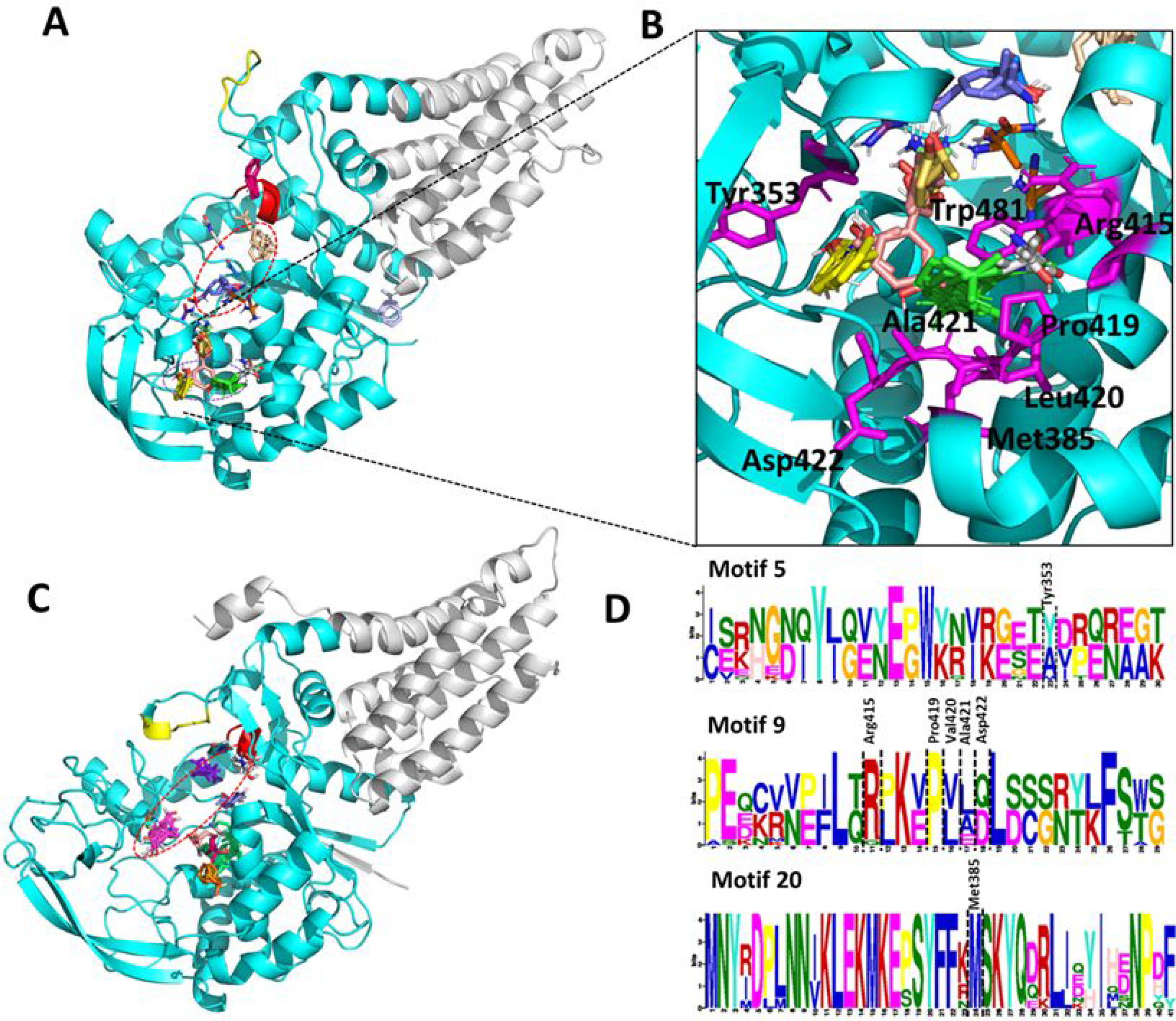
Homology models of PfMetRS and HsMetRS and prediction of potential ligand binding sites. The catalytic domain of the models is shown in cyan and the anticodon binding domain (ABD) in grey. The HIGH and KMSKS motifs are shown in red and yellow respectively. **A)** PfMetRS homology model. Known druggable sites are shown by the red dotted ellipses while the predicted site in PfMetRS by FTMap is shown in purple dotted ellipses. **B)** Insert - zoomed view of the predicted druggable pocket in PfMetRS showing stick representation (magenta) of residues interacting with probes at this site. The predicted site is located at the catalytic domain **C)** HsMetRS homology model. HsMetRS had no probable druggable sites predicted by FTMap. **D)** Motif 5, 9 and 20 logos showing conservation of residues in this family and PfMetRS residues interacting with probes predicted site.

**Figure 13:**
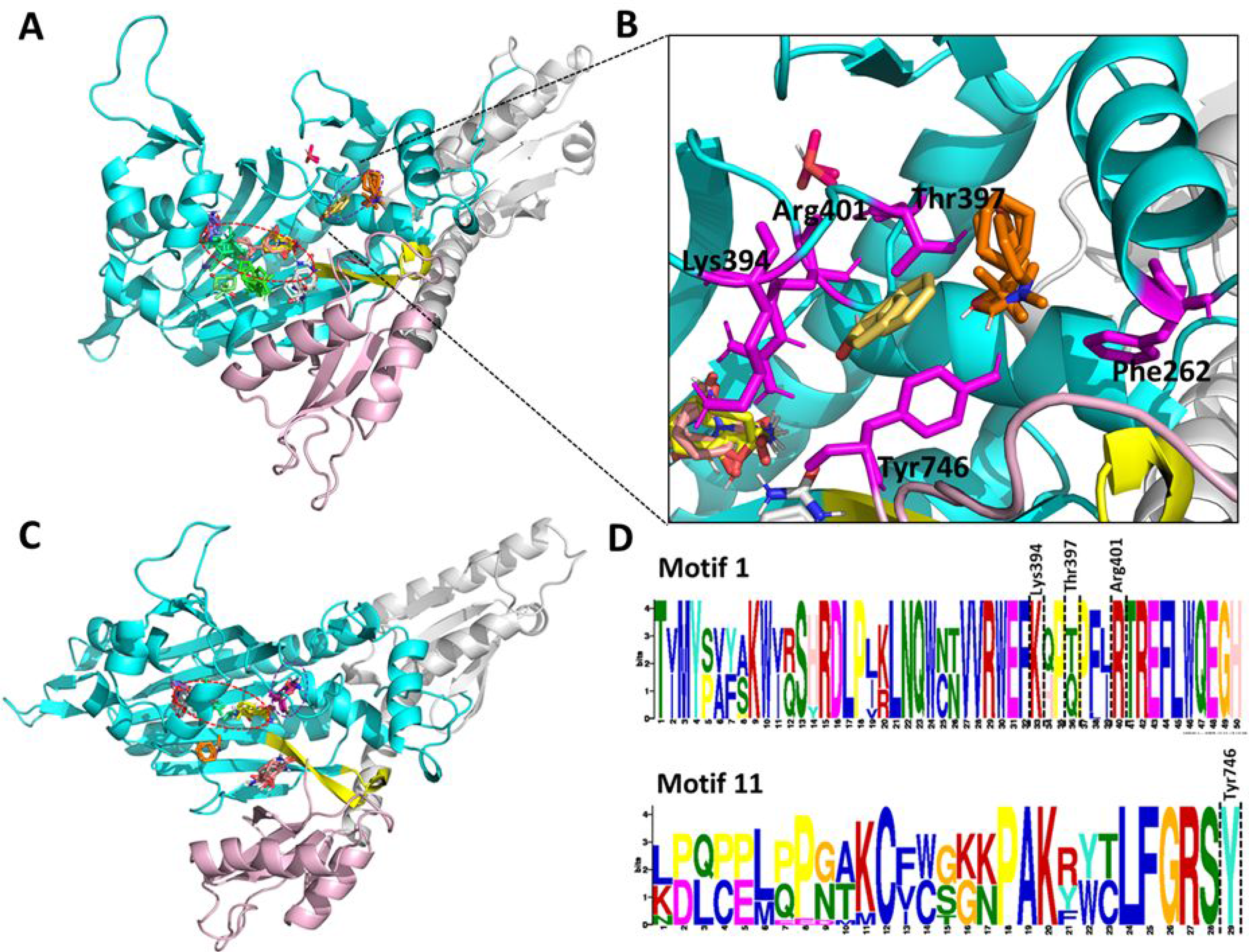
Build homology models of ProRS and prediction of potential ligand binding sites. The catalytic domain is shown in cyan, anticodon binding domain in grey and the C-terminal zinc-binding like domain is shown in light pink colour. Motif 2 located at the catalytic domain is shown in yellow. Known druggable sites are shown by the red dotted ellipses while other predicted sites by FTMap are shown in purple dotted ellipses. A) The homology model of PfProRS. B) Insert - zoomed view of the predicted site in PfProRS showing residues interacting with probes at this site as magenta sticks. These probes were interacting with residues - Tyr746, Thr397, Phe262, Arg401 and Lys394. C) The homology model of HsLysRS showing a probable druggable site with residues Thr1164, Phe1161, Thr1277, Leu1162, Thr1276 and Arg1278 interacting with probes. **D)** Motif 1 and 11 logos showing conservation of residues in these motifs and PfProRS residues interacting with probes at the predicted site.

The identified potential druggable site in PfArgRS is in a region located at the anticodon binding domain characterized by Motif 4 and 6 but the site is not present in HsArgRS (Figure 9 & 11, Table 3). Probes at this site interact with residues in the ABD - His515, Lys518, Ile522, Lys534, Glu537, Asp541 and Tyr34 located in the N-terminal domain. Motif results showed low conservation of these residues with His515 corresponding to Cys577, Lys518 to Arg580, Ile522 to Ile584, Lys534 to Thr592, Glu537 to Asp595, Asp541 to Glu599 and Tyr34 to Ser105 in the human homologue (Figure 11D). These residues are, however, highly conserved in the other plasmodium sequences studied (Additional file 4A). This region can thus be potentially targeted for inhibitor design with high selectivity to the plasmodium protein as indicated by the low conservation in the human homologue.

The predicted hotspot in PfMetRS is in a pocket formed by Motifs 5, 9, 14, 20 and the loop region of Motif 4 (Figure 10 & 12). Motif 5 is present in HsMetRS, but this motif occurs in the anticodon binding domain while Motif 14 is not present in HsMetRS (Figure 10 & 12). HsMetRS, however has a Motif 7 present in this site which is absent in the PfMetRS. Probes at the PfMetRS predicted site were interacting with residues Trp481, Ala421, Asp422, Arg415, Pro419, Met385, Leu420, Leu423 and Tyr353. Tyr353, Leu420, Asp422 and Ala421 located in Motif 4 and 9 corresponds to Ala733, Val50, Gln52 and Leu51 respectively in the human homologue. The low conservation of residues in these two motifs may explain why the probes only docked to PfMetRS and not HsMetRS. This difference in conservation at residue level in the predicted site can thus be targeted for the potential development of drugs of that bind selectively to PfMetRS. A study by Hussain *et al* [19] reported an auxiliary binding site different from ATP and methionine binding sites in PfMetRS. Inhibitors at this site interacted with residues Phe482, Ile231, His483, Tyr454, Trp447, Ile479 and Leu451 [19]. These residues map to Motif 4 and 14 located at the predicted site by FTMap in PfMetRS homology model (Table 3). An auxiliary binding pocket has also been reported in *Trypanosoma brucei* MetRS [157].

The identified potentially druggable site in PfProRS occurs at a region characterised by Motif 1, 5 and 11 which are also present in HsProRS (Figure 9 & 13). In PfProRS, residues Tyr746, Thr397, Phe262, Arg401 and Lys394 were interacting with probes docked at this site while in human, Thr1164, Phe1167, Thr1277, Leu1162, Arg1278 and Thr1276 were interacting with the probes. All residues implicated in the interaction of probes in PfProRS were conserved in all the studied sequences in this family except Thr397 which corresponds to Gln1159 in the human homologue (Figure 13D & Additional file 4R). A previous study by Hewitt *et al* [60] reported selective binding of glyburide and TCMDC-124506 at the PfProRS predicted site. This pocket is located at a region formed by α5 (residues 513-524), α9 (residues 261-272) and β-hairpin 1 and 2 (residues 276-287). We observed interaction of Phe262 to the FTMap probes and Tyr746 (Figure 13) which is also reported to interact with glyburide and TCMDC-124506 [60]. Inhibition of PfProRS by the two compounds is known to be through distortion of the ATP binding site [60]. Binding of glyburide and TCMDC-124506 causes movement of a loop between Val389 and Glu404 displacing Phe405, Arg401 and Arg390 which are key residues in ATP binding [60]. The unique predicted sites in PfArgRS, PfMetRS and PfProRS can thus be targeted through high throughput screening to identify new inhibitors.

## Conclusion

Resistance and selectivity remain a challenge when designing anti-parasitic drugs. This study aimed at getting insights on the differences at sequence and structure level between plasmodium and human aaRS. Motif analysis of the two aaRSs classes showed family specific motifs. Further, analysis of motifs for each family showed plasmodium specific and also mammalian specific motifs. Multiple sequence alignments and motif analysis of aaRS families showed high conservation of the core domains while N- and C-termini of most families showed low conservation. Interestingly, the core domain of LeuRS sequences showed low conservation despite functional conservation. ArgRS sequence alignment showed mammalian specific inserts at the N- and C-termini while mammalian TyrRS and ValRS had N-terminal extension not present in plasmodium sequences. Inhibitors can be designed to target the highly variable ABD located either at the N-terminal or the C-terminal.

On doing pairwise sequence identity calculations, ProRS was the most conserved aaRS family while GlyRS was the least conserved. Phylogenetic studies showed that human proteins had different evolutionary history to plasmodium proteins with plasmodium sequences clustering together. Plasmodium sequences also showed high sequence identity compared to the human homologues which had below 40% sequence identity. *P. yoelii* and *P. bergei* were seen to cluster in trees in most of the aaRS families showing that these proteins are closely related, and this was also depicted by the high sequence identity and shared motifs among them. *P. fragile*, *P. knowlesi* and *P. vivax* aaRSs were also seen to share evolutionary history and had high sequence identity. Prediction of additional druggable sites identified hot spots in PfArgRS, PfMetRS and PfProRS. The identified sites showed low conservation and variation of identified motifs between *P. falciparum* proteins and the human homologues. The identified sites can thus be targeted to develop drugs that only selectively bind to plasmodium proteins. As per the results in this study, it is evident that despite structural conservation, plasmodium aaRS have key features that differentiate them from human proteins. These differences can be targeted to develop antimalarial drugs with less toxicity to the host.

## Author contributions

Ö.T.B designed the study. D.W.N acquired the data, performed data analysis and wrote the initial draft. All authors contributed in interpretation and discussion of results and writing of the manuscript.

## Acknowledgements

This work is supported by the National Research Foundation (NRF) South Africa (Grant Number 105267). The content of this publication is solely the responsibility of the authors and does not necessarily represent the official views of the funders. D.W.N thanks Rhodes-Henderson Bursary and National Research Foundation South Africa for financial support. D.W.N thanks Vuyani Moses for his guidance in performing the calculations and data analysis.

## Additional files

### Additional file 1

A table showing the data set used in the study with Blast details and crystal structures retrieved from the Protein Data Bank. The species, E-value, identity, accession number, PDB ID and sequence lengths are given.

### Additional file 2

Homology model validation results obtained for Verify 3D, QMEAN and ProSA webservers. The z-DOPE scores for each model and the templates used for modelling are also shown.

### Additional file 3

Motifs discovered for the 20 aminoacyl tRNA synthetase families using MEME software. The default motif width of 6-50 residues was used. The Mast tool was used to identify overlapping motifs. The number of motifs run for each family varied and motif conservation was presented as number of sites divided by total number of class sequences and results displayed as heatmaps. Motif conservation increases from blue to red.

### Additional file 4

Results on mapping of discovered motifs on multiple sequence alignments for the 20 aaRS families. Multiple sequence alignment was performed using TCOFFEE software with default parameters.

### Additional file 5

Mapping of unique motifs to homology models in plasmodium ArgRS, MetRS, TrpRS, TyrRS, LysRS and ProRS families and the respective human homologues. Motif numbering for each protein is based on the MEME results.

### Additional file 6

Phylogenetic trees and pairwise sequence calculations for aaRS families: Molecular Phylogenetic calculations were performed using MEGA7. Sequence identity calculations were done using an in-house python script and results displayed as heatmaps. Conservation increases from blue to red.

## Abbreviations

aaRS: aminoacyl tRNA synthetases
RF: rossmann fold
CPI: connective peptide I
MSA: multiple sequence alignment
CD: catalytic domain
ABD: anticodon binding domain

